# Theft of Host Transferrin Receptor-1 by *Toxoplasma gondii* is required for infection

**DOI:** 10.1101/2023.06.23.546322

**Authors:** Stephen L. Denton, Alexa Mejia, Lindsay L. Nevarez, Miguel P. Soares, Barbara A. Fox, David J. Bzik, Jason P. Gigley

## Abstract

Nutrient acquisition by apicomplexan parasites is essential to drive their intracellular replication, yet the mechanisms that underpin essential nutrient acquisition are not defined. Using the apicomplexan model *Toxoplasma gondii*, we show that host cell proteins including the transferrin receptor 1, transferrin, ferritin heavy and light chains, and clathrin light chain are robustly taken up by tachyzoites. Tachyzoite acquisition of host cell protein was not related to host cell type or parasite virulence phenotypes. Bradyzoites possessed little capacity to acquire host cell proteins consistent with the cyst wall representing a barrier to host cell protein cargo. Increased trafficking of host cell transferrin receptor 1 and transferrin to endolysosomes boosted tachyzoite acquisition of host proteins and growth rate. Theft of host transferrin 1 and transferrin did not significantly affect iron levels in the tachyzoite. This study provides insight into essential functions associated with parasite theft of host iron sequestration and storage proteins.

## Main

The phylum Apicomplexa consists of diverse eukaryotic intracellular parasites that infect every prominent animal taxon, including humans. Malaria (*Plasmodium spp.*) ^1^, Cryptosporidiosis (*Cryptospridium spp.*) ^2^, and Toxoplasmosis (*Toxoplasma gondii*) ^3^ are common and widely distributed infections that affect a majority of the human population. These apicomplexan parasites reside in diverse host cells and acquire essential nutrients from various host cell compartments. Inside the infected host cell, *T. gondii* resides in a distinct and unique membrane enclosed structure, the parasitophorous vacuole (PV). Previous studies have identified potential pathways for host-derived nutrient transit to the rapidly replicating form of *T. gondii*, the tachyzoite, that replicates inside the PV. These mechanisms include a PV pore ^4^, the Host Organelle-Sequestering Tubulostructure (H.O.S.T.) ^5^, and the Endosomal Sorting Complex Required for Transport (ESCRT) ^6^. Recently published studies have identified the *T. gondii* micropore to provide a mechanism that sequesters host nutrients and proteins that were present in the host cell cytosol or Golgi ^7^, and to provide a mechanism that regulates homeostasis of tachyzoite plasma membrane (PM) proteins ^8^. However, previous studies have not addressed how *T. gondii* acquires the essential nutrient iron or whether *T. gondii* acquires host cargo associated with canonical mammalian host cell iron acquisition mechanisms.

Ferric and ferrous iron act as important cofactors in many cellular processes such as respiration, radical oxygen and nitrogen species production and management, gas exchange, metabolism, and transcription and translation ^9^. Because iron is involved in so many cellular processes, regulation of iron homeostasis is a critical process that is tightly controlled. Extracellular pathogens have developed many complex pathways by which they can acquire iron from the host to maintain their iron homeostasis. Their pathways can include production of bacterial siderophores that bind iron and then facilitate iron uptake by the pathogen, as well as pathogen mimicry of cellular iron transport machinery that allows the pathogen to compete against the host for available iron ^10, 11–13^. Intracellular Apicomplexan parasites also require iron for successful infection ^14, 15^. However, due to their novel location inside the host cell being surrounded by a PV, the mechanisms by which *T. gondii* acquires iron remains undefined.

Successful *T. gondii* infection is intensely dependent upon acquiring a wide variety of host nutrients ^16–18^. This demand includes iron in view that *T. gondii* growth is halted by iron chelation ^19, 20^. In enterocytes, growth inhibition by IFNγ, a major host product that controls *T. gondii* infection, is rescued by iron salt supplementation and by iron-loaded holo-transferrin ^20^. These previous studies suggest that iron restriction could be an effective mechanism to control *T. gondii* infection. *T. gondii* also possesses mechanisms that tightly regulate iron homeostasis. *T. gondii* expresses a Vacuolar Iron Transporter (VIT), protecting the parasite from iron intoxication ^19^. The requirement for iron in *T. gondii* biology includes, but is not limited to, iron-sulfur complex loading, heme biosynthesis, cytochrome cofactors, reduction/oxidation, ROS management, and a divergent respiratory complex ^21–27^. In each of these essential cellular pathways, genetic knockouts of bottleneck genes are lethal to the parasite, indicating multiple reasons why host iron is essential for parasite infection. Despite the critical importance of iron in *T. gondii* parasite biology, the mechanism(s) of iron acquisition have not been previously addressed.

In this study, we show that the genome of *T. gondii* is highly conserved for genes involved in heme-biosynthesis and iron sulfur loading complex. However, *T. gondii* is largely devoid of genes encoding intact iron storage and transport pathways. The sparse iron storage and transport gene candidates identified do not encode a complete pathway and have relatively low fitness costs when knocked out, suggesting the parasite heavily relies on host machinery and host iron. In mammalian host cells, iron transport is typically dependent on PM localized Transferrin Receptor 1 (TfR1) binding to its ligand di-ferric transferrin (holo-transferrin), followed by clathrin mediated endocytosis (CME) of the complex ^28^. Acidification of the resulting endosomes release iron from transferrin (Tf) allowing iron to be pumped into the host cell cytosol for direct use by the cell or for storage by the Ferritin heavy/light (FHC/FLC) complex ^29, 30^. Our results reveal that *T. gondii* tachyzoites directly acquire host FHC/FLC, TfR1, and Tf. The acquisition of FHC/FLC and TfR1 occurs across different host species and parasite strain types, demonstrating a highly conserved and essential host-parasite interaction. Host TfR1 was inherited by newly formed daughter tachyzoites throughout endodyogeny. Theft of host TfR1 and Tf by *T. gondii* was required for parasite growth, but unexpectedly, iron content per parasite was not dependent on sequestration of host TfR1 by tachyzoites. While host clathrin light chain (CTLC) also was ingested by tachyzoites, increased endolysosomal trafficking of TfR1 significantly boosted parasite growth as well as TfR1 ingestion. In addition, host TfR1 was not significantly present in encysted bradyzoites of *T. gondii.* Altogether, our study reveals theft of host iron transport machinery TfR1 and Tf is essential for tachyzoite replication and successful acute infection and reveals a novel mechanism that sequesters host proteins as integral components of the parasite body.

## RESULTS

### *Toxoplasma gondii* lacks conserved iron storage and transport machinery critical for iron **Homeostasis**

*T. gondii* is heavily reliant on iron for many biological processes during infection including heme biosynthesis and iron-sulfur complex loading ^22, 31^. Iron is also essential for *T. gondii* divergent respiratory chain pathway ^23, 27^. *T. gondii* is highly sensitive to its iron homeostasis, as experimentally limiting iron prevents proliferation of the parasite *in vitro* while iron intoxication also prevents proliferation ^19, 20, 32^. To dissect the molecular mechanisms involved in the use of iron and how *T. gondii* maintains iron homeostasis during infection we took an *in silico* approach to identify conserved genes in the genome of the parasite that might have a role in iron biology. Genes across all genera (plant, animal, plant, fungi, *etc*.) were organized into 5 categories: <u>1) Iron-Sulfur (Fe-S)</u> <u>Loading Complex:</u>, that facilitates irreversible loading of iron in iron-bearing proteins, <u>2) Heme</u> <u>Biosynthesis,</u> enzymes required for *de novo* heme synthesis, <u>3) Redox Functionality</u>, involved with iron-dependent functions such as ROS/RNS management and production, as well as ATP production and energy transfer in the electron transport chain, gas exchange, and membrane potential, <u>4)</u> <u>Transcription and Translation</u>, modulating the transcription or translation of any genes identified in the other categories, or proteins dependent upon direct iron signaling, and <u>5) Iron Storage and transport</u>, pathways mediating iron acquisition and sequestration. These categories of proteins were queried for protein homology in the *T.gondii* ME49 genome, then cross referenced for CRISPR-fitness scoring metadata ^33^ and ToxoDB.org (Supplementary Table 1, Figure 1A-C). The *T. gondii* genome contains complete homologs of (category 1) Fe-S Complex Loading pathways, including the ISC Pathway of bacteria, the Suf pathway of photosynthetic organisms, and the Nif pathway of eukaryotes ^26, 34^ as well as the heme biosynthesis pathway (category 2) ^22^. *T. gondii* did not have a homolog for heme catabolism proteins heme oxygenase 1 (HO-1) or pathogen-associated HemO, suggesting that the parasite may be deficient in heme salvaging. In addition, as expected from previous studies ^35–38^, redox functionality (category 3) was highly conserved in the *T. gondii* genome, which possesses several canonical iron-dependent functional proteins, that included superoxide dismutase and catalase for ROS management, aconitase as well as a ROS-sensitive divergent ATP synthase for ATP production, and cytochromes C and p450 and cognate cytochrome oxidase C for energy transfer in the electron transport chain. Our analysis also detected a single essential homolog to human eNOS/nNOS nitric oxide synthase (TGME49_219630) (Supplementary Table 1).

**Figure 1.**
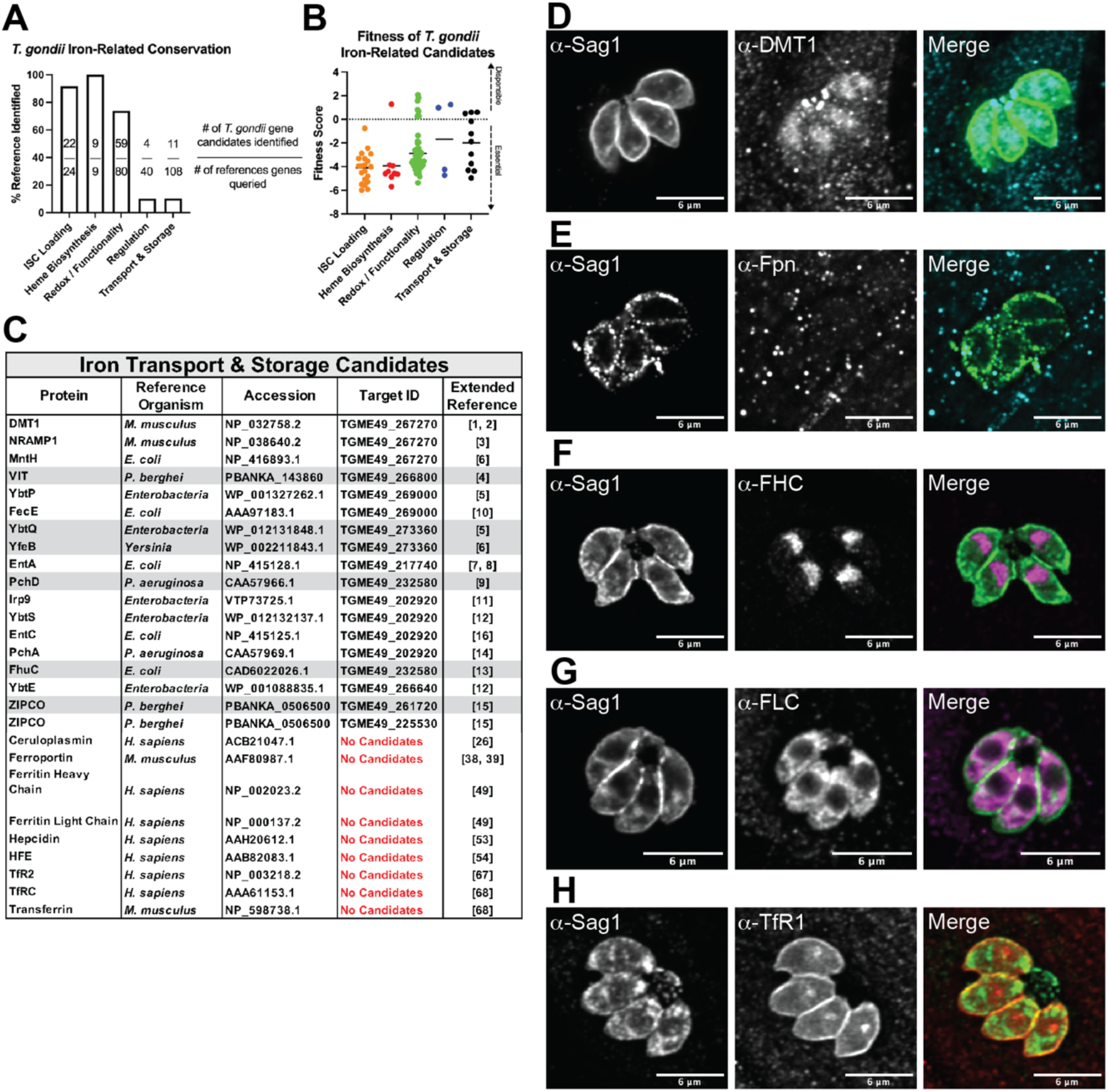
*T. gondii* lacks iron transport or storage genes but host Transferrin Receptor and Ferritin localize to the parasite. **A-C)** Known iron-related proteins were categorized according to function in 1) Iron-Sulfur Complex (ISC) Loading, 2) Heme Biosynthesis, 3) Reduction/Oxidation Production or Management (Redox / Functionality), 4) Regulation by Translation or Transcription, or 5) Storage and transport of Iron. Protein sequence data was used as query to probe the ME49 *T. gondii* genome for homology. **A)** Category fulfilment as a function of the number of *T. gondii* genes identified out of the number of proteins queried. **B)** Fitness metadata score of identified genes. **C)** List of identified genes assigned to category 5, and non-homologous mammalian category 5 genes. **D-H)** 24-hour post infection RH strain *T. gondii* confocal IFA staining for host category 5 proteins **D)** Divalent Metal Transporter 1, **E)** Ferroportin, **F)** Ferritin (Heavy Chain), **G)** Ferritin (Light Chain), and **H)** Transferrin Receptor 1.

Our bioinformatic analysis revealed that *T. gondii* expresses a very limited array of iron dependent transcription and translation (category 4) genes. We identified a single candidate with high probability to function as Iron Regulatory Protein 1 (IRP1, TGME49_226730) and two Myb-domain containing proteins, in particular BFD-1 which is the master regulator of differentiation ^39^. Intriguingly, our analysis revealed only 10.6% coverage of candidates homologous to queried iron storage and transport proteins (category 5). However, the identified genes do not establish an intact extracellular iron transport pathway or any recognizable intracellular iron storage pathway (Figure 1C). Our analysis identified four candidate parasite specific intracellular iron transporters (Figure 1C) including a transporter with homology to human DMT1 and *Escherichia coli* MntH (TGME49_267270) ^40, 41^. The other identified iron transporters were Vacuolar Iron Transporter (VIT) (TGME49_266800) ^19^, ZIPCO (TGME49_261720), and ZIPCO (TGME49_225530) ^42^. The remaining identified candidates possess homology to disparate siderophore anabolism enzymes and transporters (Figure 1C). Together, this bioinformatic analysis suggests that *T. gondii* is heavily reliant on host mechanisms of iron acquisition and host stores of iron.

### Host iron storage and transport machinery are acquired by *T. gondii* tachyzoites within the PV

Transferrin Receptor 1 (TfR1) binding to Fe^3+^-bound transferrin (Tf) on the host cell surface is the dominant ingestion pathway by which mammalian cells acquire iron from the extracellular milieu and the resulting transport of ingested TFR1-Tf associated iron to the cell cytosol is dependent upon DMT1 ^29, 43^. Ferritin Heavy (FHC) and Light (FLC) chains form a complex that can then store ingested iron as Fe^2+^ to prevent iron toxicity, but also to maintain an available intracellular iron pool ^44^. Cellular export of iron is performed by Ferroportin (Fpn) ^45^. Since *T. gondii* lacked genes with any homology to these proteins except DMT1, we investigated the association of DMT1, Fpn, FHC, FLC, and TfR1 with tachyzoites in the parasitophorous vacuole (PV). Single PV’s identified by α-Sag1 IFA (marking the tachyzoite PM) demonstrated bright α-DMT1 puncta at the apical end of the parasite (Figure 1D). However, given the high homology of parasite DMT1 to host DMT1 this result was not explored further. Host α-Fpn exhibited no specific association with the PV and no apparent change of localization in infected cells (Figure 1E). In contrast, IFA demonstrated intense host α-FHC, α-FLC, and α-TfR1 staining at the parasite location (Figure 1F-1H).

Based on localization of fluorescence in reference to α-Sag1 staining, the sequestration of TfR1 appeared to outline the periphery of the parasite PM and/or inner membrane complex (IMC) in a complete encasement, with a singular sequestered spot of TfR1 fluorescence (Figure 1H, 2A-C). On occasion, TfR1 could be observed outside individual parasites but within the PV lumen, perhaps representative of a chance observation of TfR1 *en route* to the parasite (Figure S1C). Notably, the mean fluorescent intensity (MFI) of TfR1 in the PV was significantly higher than the surrounding host cells (Figure 2D-F). The presence of TfR1 was confirmed by western blotting of purified parasite lysate compared to uninfected host cell lysate (Figure 2G, Figure S2). The identified apparent molecular weight of the ∼97kDa TfR1 band was slightly downshifted in purified parasite lysate, possibly indicating processing events occurred during or after TfR1 theft. This possibility may also explain different staining patterns of TfR1 in the parasite when using different antibody clones (Figure S1A and S1B). The cargo of TfR1, Tf, also localized in the parasite and had a higher MFI in the PV than surrounding host cells (Figure 2H-2I, and Figure S1D). This Tf localization phenotype displayed a less prominent peripheral localization and a more distinctive punctate pattern than TfR1.

**Figure 2.**
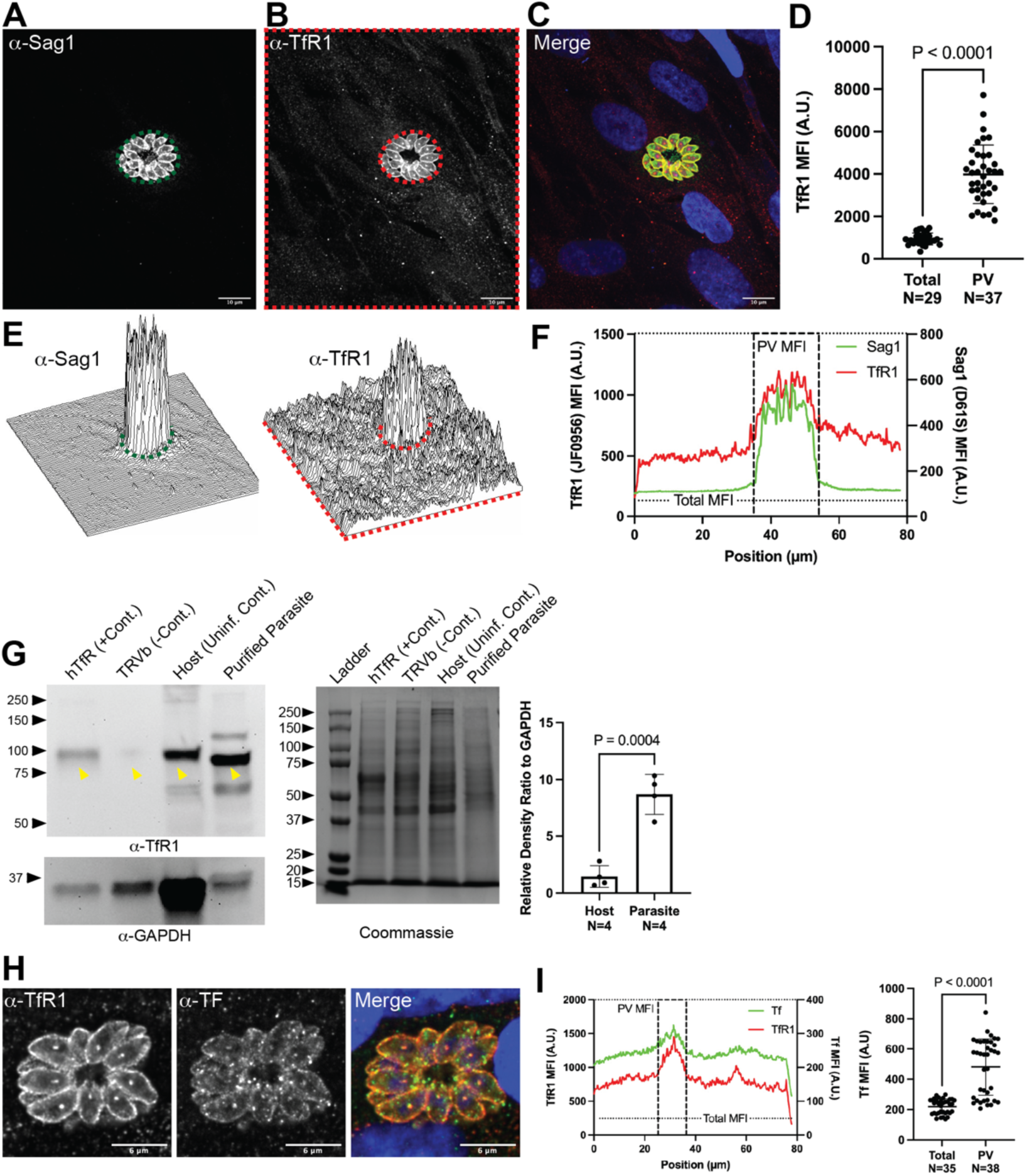
Host transferrin receptor 1 and cognate ligand transferrin are acquired by *T. gondii*. **A-F)** 24-hour post infection RH strain *T. gondii* and surrounding uninfected cell confocal IFA staining for host transferrin receptor 1 at low magnification. **A)** Representative image of α-Sag1 fluorescence to demonstrate parasite location. Parasite boundary is restricted to area digitally denoted by dashed green line. **B)** Representative image of α-TfR1 (JF0956) fluorescence. Boundary distinctions of parasite area (PV), as determined by α-Sag1 Fluorescence, and the whole image (Total) are denoted in dashed red lines. **C)** Representative image of digitally merged Sag1 (green) and TfR1 (red) and DAPI (blue) fluorescence. **D)** Comparison of relative Mean Fluorescence Intensity (MFI) of TfR1 corresponding to total image area (Total) and parasite-restricted area (PV). Data pooled from four technical repeats and analyzed by unpaired student’s t-test (t=11.70, df=64). **E)** Representative surface plots displaying pixel intensity of Sag1 (left) and TfR1 (right) as further demonstration of boundary establishment. **F)** Representative colocalization as overlayed distribution plots of y-slice MFI of Sag1 (green) and TfR1 (red). **G)** 30µg of cell lysates were probed in Western Blot for TfR1 (H68.4) (left). Positive control hTfR cell lysates are TfR1-deficient cells reconstituted with the human TfR1 gene. Negative control TRVb cells are the TfR1-deficient cell. The uninfected control is host cell lysate from uninfected MRC5 cells. Purified parasites are RH strain *T. gondii* mechanically lysed from host cells and filter purified. Coommassie stain of gel provided as additional loading control (middle). Arrows indicate area tested for densitometry (right). Graphical data pooled from four technical repeats and analyzed by unpaired student’s t-test (t=7.143, df=6). Unaltered western blot images and additional replicates are shown in Extended Data 1. **H)** High magnification representative images of parasite confocal IFA staining for host TfR1 and Tf. **I)** Representative distribution plot of TfR1 and Tf y-slice MFI; quantitation of parasite-associated Tf Quantitation pooled from three technical repeats and analyzed by student’s t-test (t=8.028, df=71).

Single PVs also demonstrated significantly increased FHC and FLC MFI compared to surrounding host cells. FHC staining in tachyzoites appeared to be centrally localized the parasite whereas FLC appeared to be throughout the cytoplasm of *T. gondii* (Figure 1F-G, Figure S3A and S3B). Whether FHC and FLC colocalized with each other could not be determined.

Differential permeabilization allows differential IFA penetration to the parasite cytosol, parasite surface and PV lumen, or PV inner leaflet using Triton X-100 (as in Figures 1 and 2) ^46^, Saponin ^47^, and Digitonin ^48^, respectively. To more precisely assess the localization of TfR1, FHC, and FLC, we performed IFA following each permeabilization method, and observed that the parasite morphology of TfR1, FHC, and FLC were occluded after Saponin or Digitonin permeabilization (Figure 3A and Figure S4A-S4B). Regardless of permeabilization method, the MFI of TfR1 and FLC was increased at the parasite location compared to the surrounding area (Figure 3B and Figure S3F). However, the ratio of increase of parasite associated TfR1, FHC, and FLC were each significantly greater in Triton X-100 permeabilized samples compared to Saponin or Digitonin permeabilized samples (Figure 3C, Figure S4C and S4E). These results confirm that TfR1, FHC, and FLC localize and accumulate within replicating PV-localized tachyzoites.

**Figure 3.**
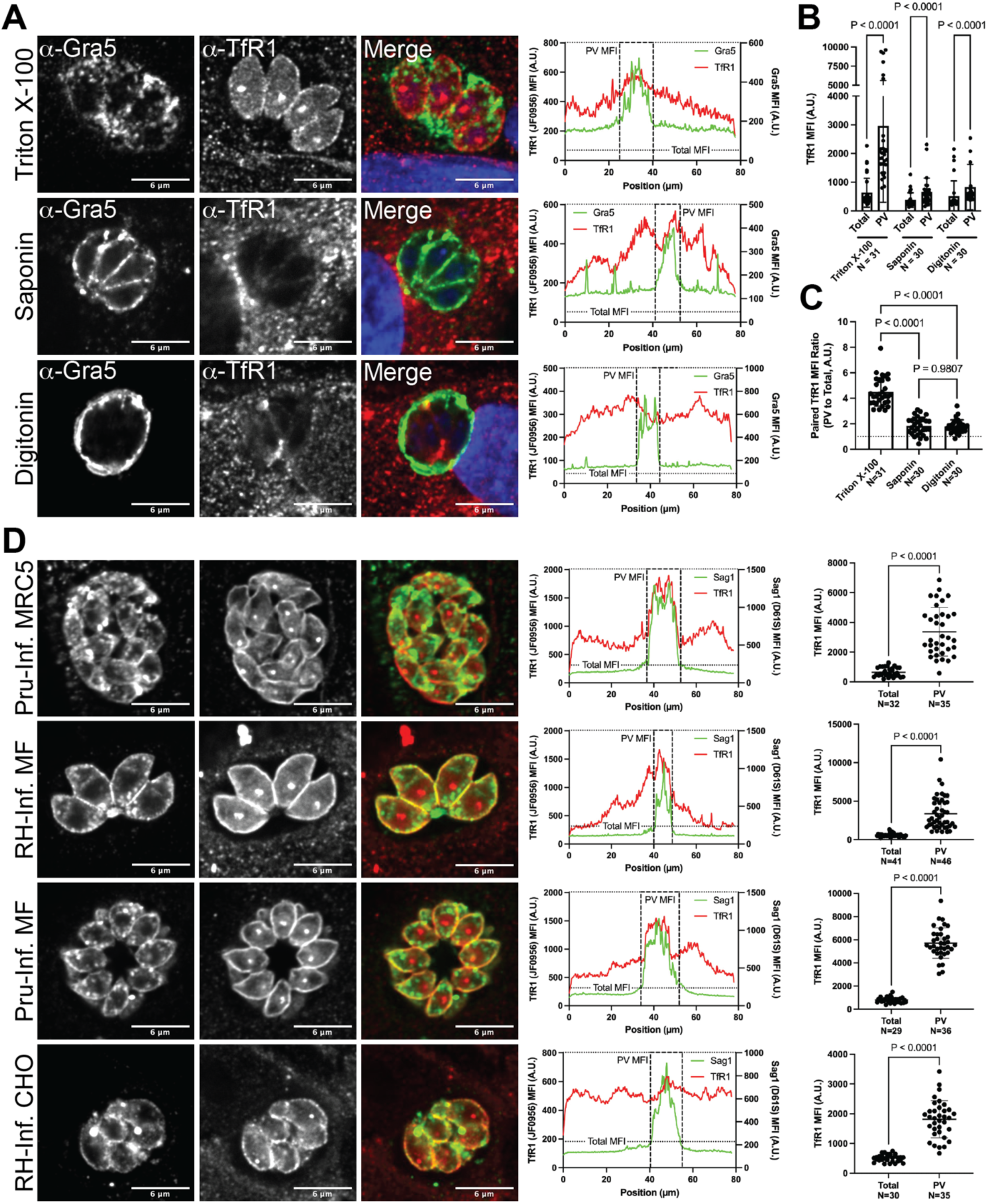
Transferrin receptor 1 is incorporated into parasite cells irrespective of virulence type or host cell genus. **A-C)** RH-strain *T. gondii* 24-hour infections of MRC5 cells were differentially permeabilized and stained for confocal IFA of host TfR1. **A)** Shown are representative confocal images for Sag1, TfR1, and merged fluorescence and corresponding distribution plots for *Top Row:* Triton X-100 permeabilized cells, *Middle Row:* Saponin permeabilized cells, and *Bottom Row:* Digitonin permeabilized cells. **B)** Comparisons of TfR1 MFI in Total image area and parasite-associated PV area, grouped by permeabilization type. Data pooled from two technical repeats and analyzed by Two-Way ANOVA (f=17.91, df=178). **C)** Comparison of TfR1 PV/Total ratio between permeabilization type. Data pooled from two technical repeats and analyzed by Ordinary One-Way ANOVA (f=111.5, df=90). **D)** RH-strain or Prugniaud (Pru) strain *T. gondii* inoculated into MRC5, MF, or CHO host cells were assayed for confocal TfR1 IFA. *First (Top) Row:* Representative images, distribution plot, and quantitation of parasite associated TfR1 of Pru-strain *T. gondii* infecting human MRC5 cells. *Second Row:* Representative images, distribution plot, and quantitation of parasite associated TfR1 of RH-strain *T. gondii* infecting mouse fibroblast (MF) cells. *Third Row:* Representative images, distribution plot, and quantitation of parasite associated TfR1 of Pru-strain *T. gondii* infecting MF cells. *Last (Bottom) Row:* Representative images, distribution plot, and quantitation of parasite associated TfR1 of RH-strain *T. gondii* infecting hamster CHO cells. Data pooled from two independent repeats and analyzed by unpaired student’s t-test (first row, t=9.223, df=65; second row, t=8.543, df=85; third row, t=19.92, df=63; fourth row, t=11.25, df=63).

There is only one genus and species of *T. gondii*, however there are 3 primary canonical types of parasite based on polymorphisms in the genome that affect virulence in mice. Type I strains are highly virulent, and Type II and III strains are avirulent. All types of *T. gondii* can infect virtually any nucleated cell of any animal ^3^. Therefore, we tested how conserved theft of host iron storage and transport machinery was based on parasite virulence and host genera. Theft of host iron machinery was conserved for TfR1 in murine fibroblasts (MF’s) and Chinese Hamster Ovary (CHO) cells by both Type I (RH) and Type II (Pru) parasites (Figure 3D). Similarly, the phenotype of FHC/FLC theft was conserved by parasite virulence type and host cell genera (Figure S3C-S3F). Altogether, these results indicate that host iron storage and transport machinery is sequestered by *T. gondii* cells in a conserved pathway.

Given the unexpected nature of the theft of host iron related machinery, we retroactively searched for evidence of this phenomenon in other infections. *Plasmodium*, another apicomplexan parasite, is also heavily reliant on iron. Before the advent of next-generation sequencing, *Plasmodium* extract was found to bind TfR1 antibodies as well as Tf protein, demonstrating evidence of a parasite-encoded Transferrin Receptor capable of binding transferrin ^49^. Therefore, we assessed the extent of *Plasmodium* homology to human iron-related proteins. However, similar to *T. gondii*, we were unable to find homology to human iron transport or storage proteins in *Plasmodium* other than DMT1, (Table S2).

### Theft of Transferrin Receptor is inherited by daughter cells

*Toxoplasma* asexual replication occurs through a process known as endodyogeny, or “formation of two within” ^50^. We tested the fate of host TfR1 from parent cell to offspring using IFA. Tachyzoites undergoing endodyogeny were observed using staining for α-Sag1 with 10-25% of vacuoles having atypically large tachyzoites (Figure 4B). Endodyogeny was confirmed using DAPI dye where parasites were classified as replicating due to observation of two segregated nuclei confined within one parental parasite (Figure 4A). Surprisingly, the TfR1 immunofluorescence localized to the periphery of the parent tachyzoite, but also lining the periphery of the two offspring cells inside (Figure 4C). These results phenocopied the distribution of IMC1 during endodyogeny ^51^. The characteristic TfR1 puncta was also observed within the daughter cells. Host TfR1 association with newly forming offspring cells closely mirror each step of endodyogeny (Figure 4E-4J), including the apparent formation of basal buds (Figure 4F), descending IMC (Figure 4G), extension to the apical ends (Figure 4H), and breakdown of mother cell (Figure 4I). These observations further support the idea that theft of host TfR1 has a functional purpose for *T. gondii* infection.

**Figure 4.**
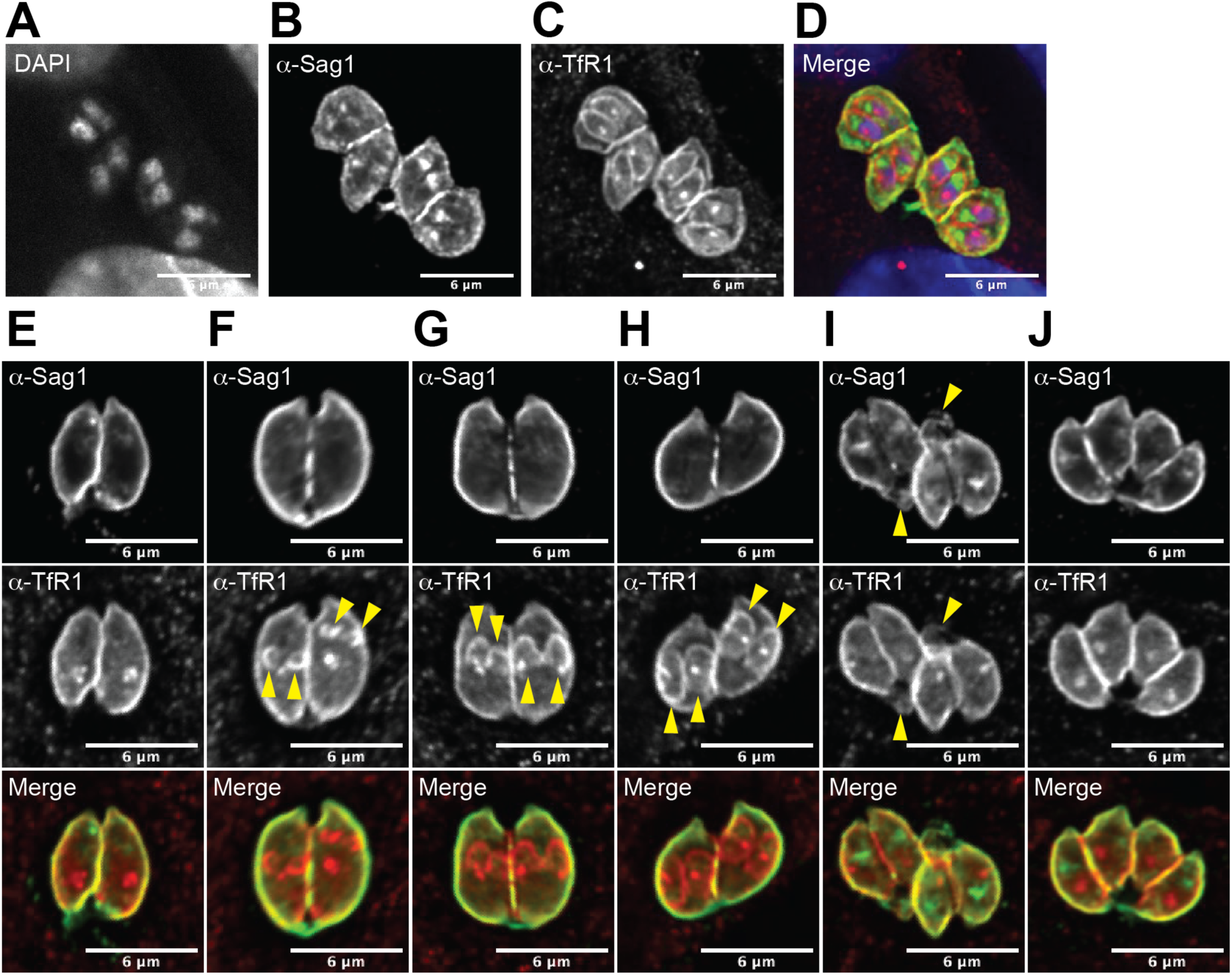
Host transferrin receptor is inherited by daughter cells during endodyogeny. RH-strain *T. gondii* infecting MRC5 cells for 15-24 hours were permeabilized with Triton X-100 and stained for confocal Sag1 and TfR1 IFA. The following phenotypes were partially penetrant. **A-D)** Exemplar images are shown for parasites undergoing endodyogeny by DAPI (A), Sag1 (B), TfR1 (C) and merged (D) fluorescence. Additional exemplar images are shown for: **E)** Pre-endodyogeny, **F)** Early Endodyogeny, arrows indicate morphology associated with initiation of scaffold assembly, **G)** Mid-endodyogeny, arrows indicate morphology associated with scaffold elongation, **H)** Late endodyogeny, arrows indicate morphology associated with organelle segregation, **I)** Cytokinesis, arrows indicate morphology associated with mother cell remnants, and **J)** Post-endodyogeny.

### Tachyzoite replication is associated with functional TfR1

The primary mechanism of TfR1-holo-Tf iron transport is highly conserved across most eukaryotes. Therefore, we focused our efforts on testing how TfR1 and holo-Tf impacted the ability to *T. gondii* to infect and replicate in host cells. Ubiqutin Cre-ER^T2^ X TfR1 (flox/flox) mice were generated as a source of fibroblasts for *in vitro* parasite proliferation assays and IFA for TfR1 (Figure 5). TfR1 mRNA expression is efficiently ablated only in Cre-possessing and tamoxifen treated cells (Figure 5A), however, at 6 days post infection there was no significant difference in parasite genomes by quantitative PCR in the TfR1 knockdown group compared to all controls (Figure 5B). The ratio of 1, 2, 4, 8 and 16 tachyzoite containing PV’s was assayed at 24 hours post infection (Figure 5C and 5D). In Cre-cells +/- tamoxifen and Cre+ cells without tamoxifen cultures there was no change in the ratios of tachyzoites per PV and similar numbers of 1, 2, 4, 8 and 16 stage PVs were quantified. Although there appeared to be a slight shift in the ratio of tachyzoites per PV in the Cre+ cells with tamoxifen TfR1 knockout cultures, the differences were modest (Figure 5D). IFA for TfR1 at 24 hours demonstrated that the parasite was still sequestering TfR1 efficiently as no significant difference in PV MFI between the different culture conditions was observed (Figure 5E). These results suggest that nascent TfR1 protein expression by the host cell is not required by the parasite for successful infection.

**Figure 5.**
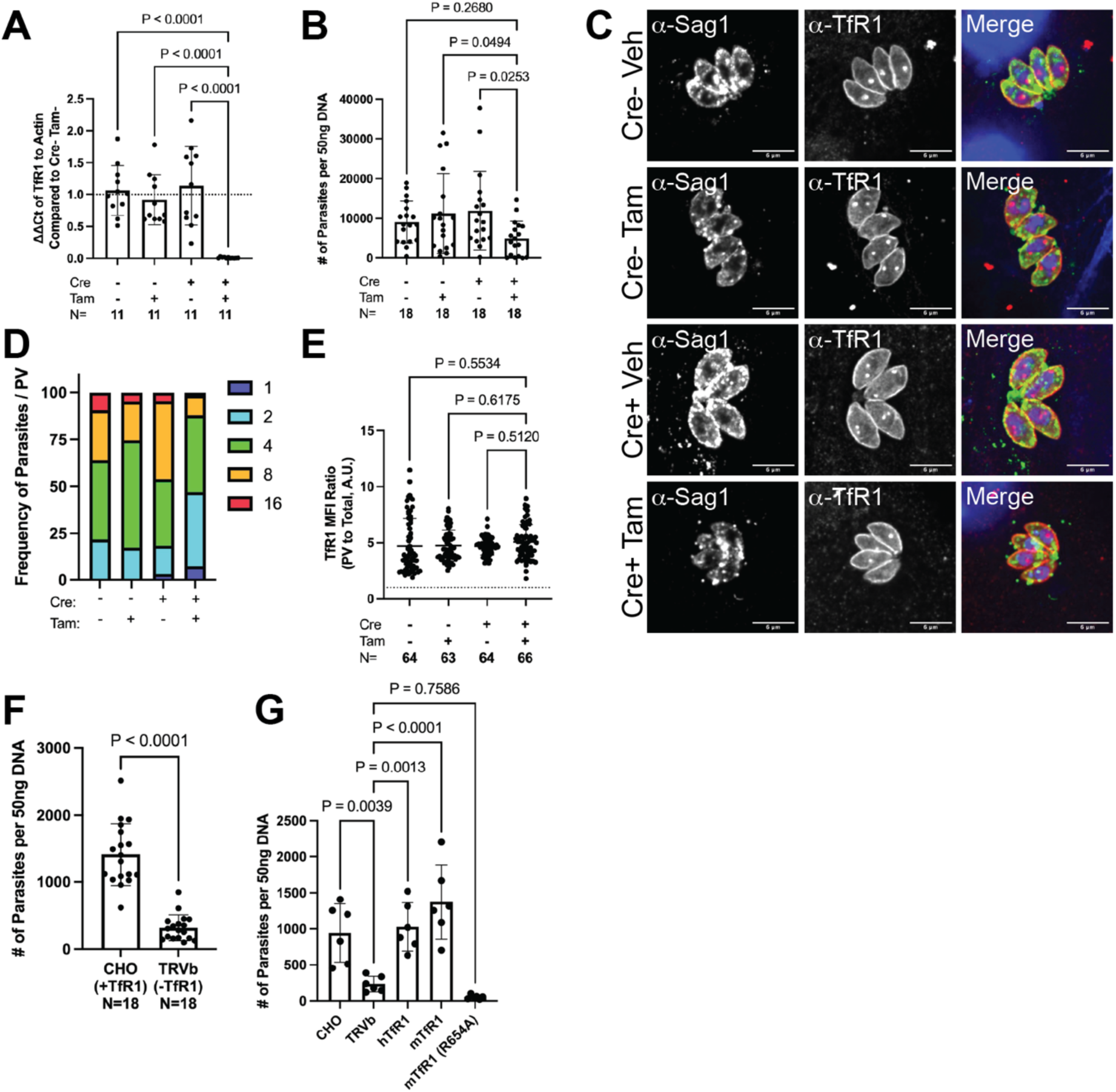
Functional TfR1 is required for successful *T. gondii* infection. TfR1^Flx/Flx^ mice were crossed with Ubiqiutin-CreER^T2^ background. Fibroblasts from these mice containing or lacking the Cre transgene were generated. **A)** Impact of tamoxifen on Cre-possessing or Cre-lacking fibroblast expression of TfR1 mRNA by quantitative reverse transcriptase PCR. Data pooled from two technical repeats and analyzed by Ordinary One-Way ANOVA (f=17.51, df=43) **B)** Impact of tamoxifen on *T. gondii* replication in Cre-possessing or Cre-lacking fibroblasts by genome equivalence in RT-PCR. Data pooled from three technical repeats and analyzed by Ordinary One-Way ANOVA (f=2.899, df=71) **C)** Representative Sag1, TfR1, and merged IFA images of groups as in A-B at 24 hours post infection. **D)** Graph shows distribution of PV’s containing 1, 2, 4, 8, or 16 parasites of groups depicted in C. Pooled proportions of two technical repeats of N=4 each is shown. **E**) Quantitation of TfR1 MFI of parasite area compared to the TfR1 MFI of surrounding total area of the groups depicted in C. Data pooled from two technical repeats and analyzed by Ordinary One-Way ANOVA (f=0.8677, df=256) **F)** TfR1-deficient TRVb cells and parental CHO cells were infected with RH-strain *T. gondii* and assessed for parasite burden. Data pooled from three technical repeats and analyzed by unpaired student’s t-test (t=9.228, df=34) **G)** Parental CHO and TfR1-deficient TRVb cells complemented with the human TfR1 gene (hTfR1), murine TfR1 gene (mTfR1), or Tf-binding site mutated murine TfR1 allele (R654A) were infected with RH and assessed for parasite burden as in F. Data representative of three technical repeats and analyzed by Ordinary One-Way ANOVA (f=16.96, df=29, N=6).

To elucidate the functional requirement of TfR1 and Tf for *T. gondii* infection we obtained TfR1 deficient Chinese Hamster Ovary (CHO) cells (CHO-TRVb1), CHO-TRVb1 cells complemented with human TfR1, murine TfR1, and murine TfR1 with a R654A point mutation that is unable to bind holo-Tf ^52, 53^ and measured *T. gondii* proliferation *in vitro*. Tachyzoite proliferation was significantly inhibited in the TfR1-deficient CHO-TRVb1 cells (Figure 5F) compared to wildtype CHO cells. Overexpression of human and murine TfR1 in the CHO-TRVb1 cells restored *T. gondii* proliferation (Figure 5G). Additionally, the R654A mutated allele was insufficient to rescue parasite growth to wild-type CHO cell levels (Figure 5G).

Internalization of TfR1 and Tf by mammalian cells is dependent upon clathrin mediated endocytosis (CME) ^28^. IFA staining for human clathrin light chain (CTLC) exhibited clear sequestration within the parasite (Figure 6A-C). Though *T. gondii* does possess a CTLC homolog, TGME49_290950, protein alignment of this gene demonstrates incompatibility with the human α-CTLC clone (2E5, Figure 6D and 6E). Another possible source of TfR1 and Tf for the parasite is the endosomal recycling complex (ERC) ^54^. The tubule extension of the ERC was unable to be visualized by EHD1/MICAL-1 IFA near the parasites nor was there any association of these molecules with parasites (data not shown), so it is uncertain if the ERC pathway is manipulated for TfR1-Tf theft. To functionally test the importance of the source of TfR1 infection we utilized A24 monoclonal antibody blockade. TfR1 protein can be reduced using the clone A24 anti-human TfR1 monoclonal antibody therapy, which enhances TfR1 CME and targets TfR1 degradation in the lysosome ^55^. Unexpectedly, treatment of human fibroblast cultures with A24 antibody compared to vehicle alone resulted in a nearly 4-fold increase (NT: 7245 +/- 5494, A24: 25184 +/- 17139) in parasite genomes detected over the non-treated control (Figure 6I). Despite higher parasite numbers in A24 cultures, TfR1 MFI in tachyzoites were not different from vehicle alone controls (Figure 6F-H). These results demonstrate that misdirecting TfR1 from the ERC to lysosome enhanced parasite proliferation and acquisition of TfR1.

**Figure 6.**
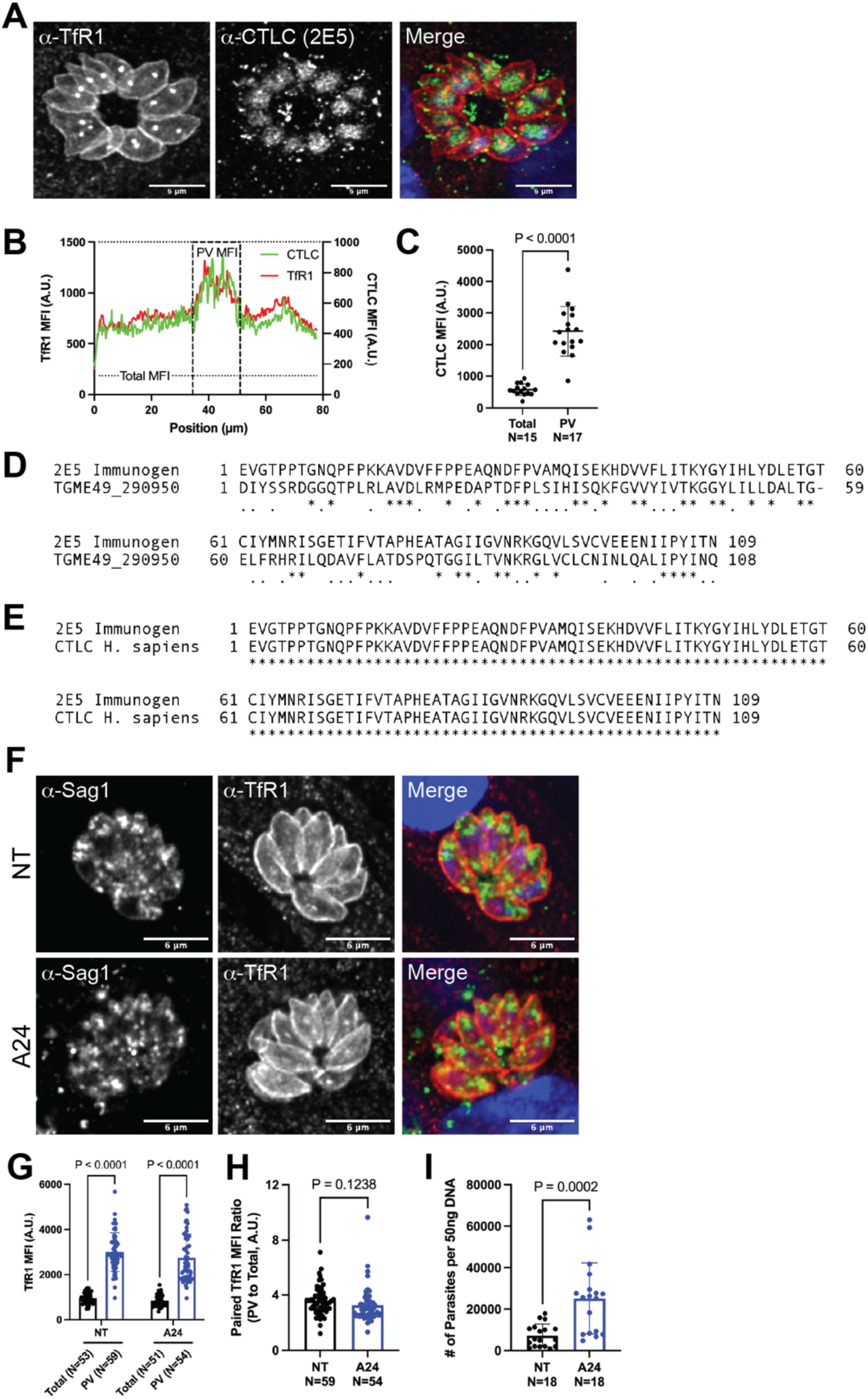
Clathrin light chain associates with *T. gondii* and blocking endosome recycling confers advantage to parasite replication. **A-C)** RH-strain *T. gondii* infecting MRC5 cells for 24 hours were permeabilized with Triton X-100 and stained for confocal Sag1 and CTLC IFA. Representative images, distribution plot, and quantitation of parasite association with CTLC are shown. Data for CTLC quantitation is pooled from two technical repeats and analyzed with unpaired student’s t-test (t=8.922, df=30) **D)** Displayed is the protein sequence alignment of the epitope used to generate the CTLC clone with the top scoring homologous *Toxoplasma* gene (TGME49_290950). **E)** Displayed is the protein sequence alignment of the epitope used to generate the CTLC clone with the human CTLC gene. **F-I)** Human MRC5 fibroblasts were treated with TfR1-ERC blocking antibody (A24) and infected with RH strain *T. gondii*. **F)** Representative images are shown of 24-hour post infection IFA for Sag1 and TfR1. **G)** Comparisons of Total- and PV-associated IFA of TfR1 MFI are shown grouped by treatment. Data pooled from two technical repeats and analyzed by Two-Way ANOVA (f=3.561, df=213) **H)** Comparison of PV/Total TfR1 MFI ratio between nontreated and A24 treated groups is shown. Data pooled from two technical repeats and analyzed by unpaired student’s t-test (t=1.551, df=111). **I)** Nontreated and A24 treated MRC5 cells were infected with RH strain *T. gondii* and assessed for parasite burden by genome equivalence. Data pooled from three technical repeats and analyzed by unpaired student’s t-test (t=4.229, df=34).

These data altogether demonstrate that acquisition of functional host TfR1 and Tf by *T. gondii* is required for successful infection.

### Theft of TfR1 is dependent on parasite life stage

After induction of the immune response, or nutritional or environmental stress, tachyzoites convert into slow growing bradyzoite persistent stages that are encased in a thick-walled cyst structure ^39, 56^. Since we observe that TfR1 theft also occurs in a cyst forming parasite strain, we tested whether bradyzoites contained within cysts in the brains of chronically infected mice also harbored host TfR1. Cysts of infected mice were identified within 40µm sections of brain tissue with FITC-conjugated *Dolichos biflorus* agglutinin lection (Figure 7A). While TfR1 was readily observed on neuronal cells in the brain section, the characteristic accumulation of host-TfR1 to parasites was not observed within the confines of the cysts (Figure 7B). At times, the TfR1 staining appeared to border the cyst, but this result was inconsistent (Figure 7B, 7D). The MFI of TfR1 of cysts was not significantly different than the surrounding area of the brain section or over background signal suggesting that TfR1 was not maintained by the bradyzoite (Figure 7E). Live Gra5+ bradyzoites released from purified and trypsinized cysts were analyzed for α-TfR1 binding by flow cytometry (Figure 7F and 7G). TfR1 signal was significantly lower in purified (non-reactivated) bradyzoites, compared to tachyzoite controls (Figure 7H). TfR1 signal in the bradyzoite population was heterogenous, however, as the MFI of TfR1 was distinctly split into two populations (Figure 7G). One population possessed TfR1 at a similar extent as *in vitro* tachyzoites, and one population exhibited no TfR1 as the fluorescence was indistinguishable from the FMO negative control. In separate experiments, the bradyzoites were reactivated and TfR1 theft occurred by ME49 strain parasites as robustly as in previous experiments (Figure 7I and 7J). These results indicate that the cyst wall is a barrier to TfR1 sequestration by bradyzoites.

**Figure 7.**
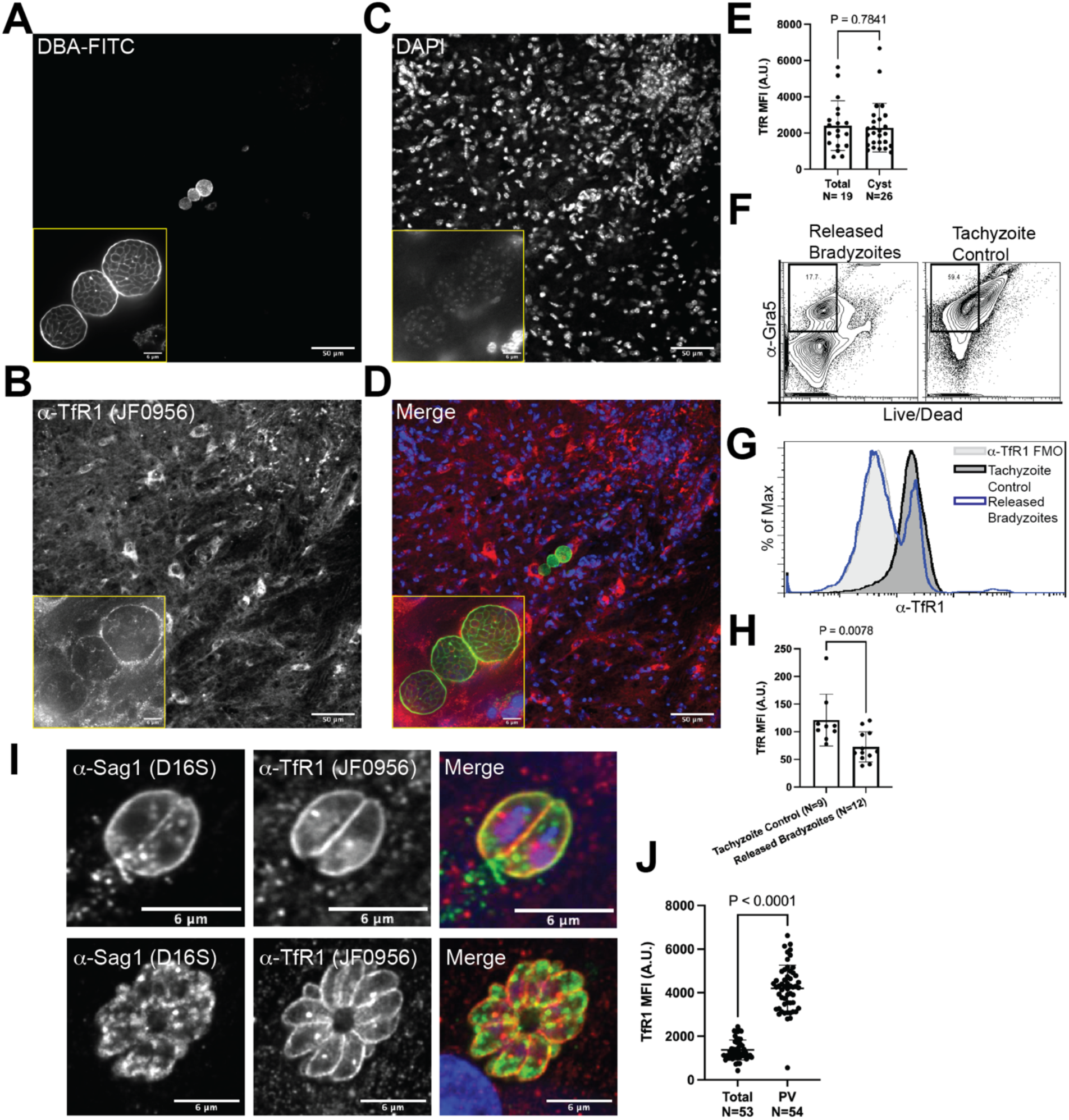
Theft of transferrin receptor 1 is dependent on life stage. **A-D)** Chronically ME49-strain *T. gondii* infected C57BL/6 brains were sectioned for IFA staining of Dolichos Bifluorus Agglutininn (DBA) and TfR1. Representative low magnification and inlaid high magnification images are shown for DBA (A), TfR1 (B), DAPI (C), and merged (D) fluorescence. **E)** Graph is shown depicting total field or cyst-restricted (boundary of DBA fluorescence) TfR1 MFI. Data pooled from two independent repeats and analyzed by unpaired student’s t-test (t=0.2758, df=43). **F-J)** Cysts purified from chronically ME49 infected mice were trypsinized and assessed for protein expression by flow cytometry or reactivated *in vitro.* **F)** Displayed are representative dotplots depicting Gra5 expression and viability (Live/Dead negative) of released bradyzoites from cysts or RH tachyzoites grown *in vitro.* **G)** Representative histogram of TfR1 protein level on Live Gra5+ parasites is shown. **H)** Relative TfR1 MFI on Live Gra5+ parasites from the released bradyzoite population or the *in vitro* tachyzoite population is compared. Data pooled from two technical repeats and analyzed by unpaired student’s t-test (t=2.975, df=19) **I)** Released ME49 bradyzoites were reactivated in human MRC5 cells for 24 hours and representative images of Sag1, TfR1, and merged fluorescence are shown. **J)** Quantitation of the parasite associated TfR1 of reactivated bradyzoites. Data pooled from two technical repeats and analyzed by unpaired student’s t-test (t=17.71, df=105).

### Theft of host TfR1 does not impact iron homeostasis in tachyzoites of *T. gondii*

We speculated that tachyzoite sequestration of host iron metabolism proteins could be a mechanism for parasite iron acquisition. To assess the extent to which the Tf-TfR1 complex is a courier of iron to the parasite, we quantified the elemental mass content of *T. gondii* on a per parasite basis by Inductively-Coupled Plasma Mass Spectometry (ICPMS). As a proof of concept, variable parasite titers of RH-strain *T. gondii* harvested from routine MRC5 culture were assessed for iron, copper, manganese, and zinc content (Figure S5). The relationship between the parts-per-billion (ppb) of each the metals correlated linearly with the parasite number. While the metals were at significantly higher levels than the limit of detection (LOD), with the exception of manganese, they were at extremely minute concentrations per parasite. Subsequently, 5*10^6^ RH-strain *T. gondii* tachyzoites each were harvested from CHO, TRVb, hTfR, mTfR, and R654A infections for ICPMS. As an internal control, quantities of elemental phosphorous and sulfur were assessed; there was no significant differences amongst the groups, indicating that the cell numbers were accurately normalized between groups (Figure 8). Amongst the groups, there was additionally no significant variation of magnesium, copper, or of elemental iron. Curiously, there was a significantly higher amount of detected zinc of parasites grown within TRVb compared to all other groups (Figure 8).

**Figure 8:**
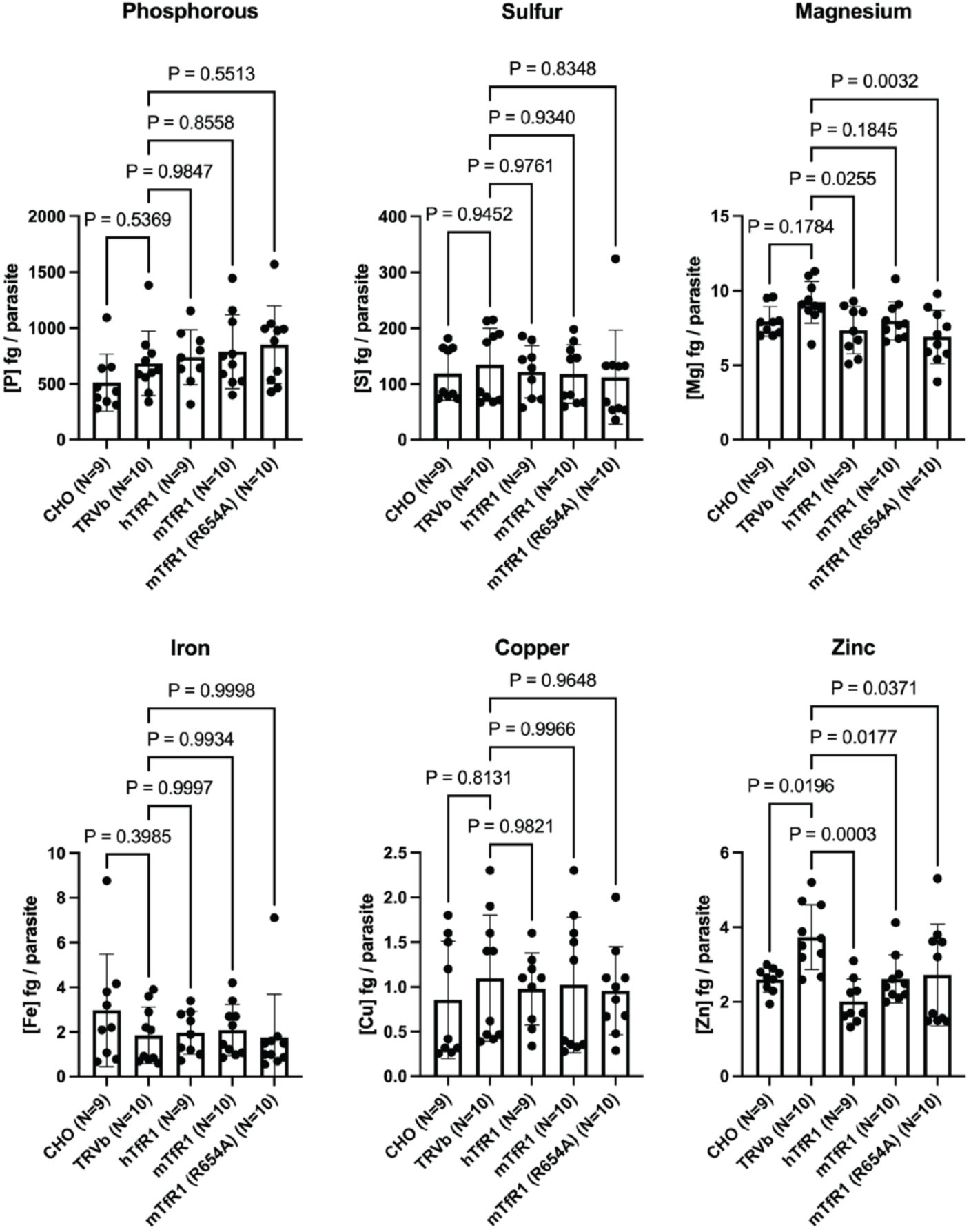
Theft of host TfR1 does not impact iron homeostasis in tachyzoites of *T. gondii.* 10^5^ Tachyzoites RH strain *T. gondii* were inoculated into CHO, TRVb, hTfR1, mTfR1, and R654A cells for 4 days and isolated from the host cells. Five million purified parasites were digested for elemental composition analysis by ICPMS. Depicted are the mass of each element on a per cell basis. Data is pooled from two technical repeats and analyzed by Ordinary One-Way ANOVA (Phosphorous, f=1.741, df=47; Sulfur, f=0.1832, df=47; Magnesium, f=3.612, df=47, Copper, f=0.1952, df=47; Zinc, f=5.212, df=47; Iron, f=0.8040, df=47).

**Figure 9.**
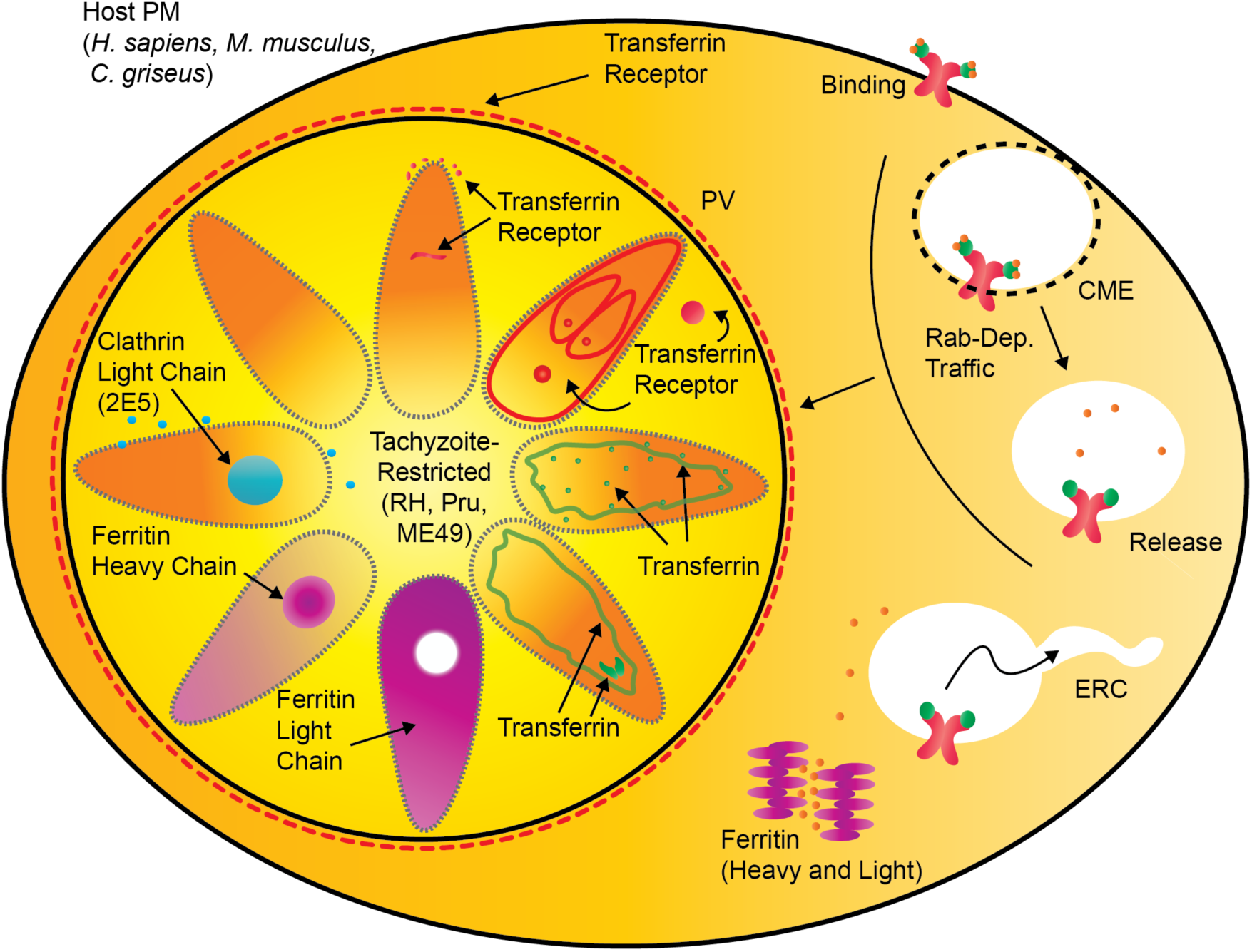
Illustrated Model: Theft of host iron machinery by *Toxoplasma gondii*. Transferrin receptor 1 (depicted in red) was found to associate near the PVM, around the periphery of the parasite cell, disperse throughout the parasite, in isolated puncta or lobules within the parasite, and deposited on growing daughter cells in endodyogeny. Transferrin (depicted in green) was also found to associate near the periphery of the parasite surface, in disperse puncta throughout the host cell, and in lobules in the parasite. Host CTLC (depicted in cyan) was detected in parasites. Ferritin heavy and light chain (depicted in magenta) were detected in parasites. Both ferritin chains and TfR1 were confirmed to incorporate within the parasite. TfR1 and Ferritin theft occurred in multiple strains (RH, Pru, ME49) and during infection of multiple host genera (*Mus musculus, H. sapiens, C. griseus)*. Host traffic of TfR1 prior to entry of endosomal recycling complex (ERC) is advantageous for the parasite.

## DISCUSSION

The protozoan parasite *T. gondii* relies on several host derived nutrients for successful infection. Iron is one of those nutrients and the parasite needs to tightly regulate iron homeostasis. The mechanisms involved in regulating iron homeostasis in *T. gondii* are not known. Here we report that *T. gondii* and *Plasmodium spp.* lack genes required for iron storage and transport. FHC and FLC form a complex that allows cells to store iron and TfR1 binding to holo-Tf is what allows cells to transport iron from extracellular sources. These proteins are critical for most mammalian cells to regulate iron homeostasis. Interestingly, *T. gondii* appears to acquire host FHC, FLC, TfR1, and Tf proteins during acute infection. This process is highly conserved across parasite virulence types and host cell genus. *T. gondii* requires functional TfR1 that can bind Tf for successful infection. Although we find evidence of host CTLC in the tachyzoites, suggesting that clathrin mediated endocytosis might be the pathway hijacked by *T. gondii* to acquires these proteins, targeting TfR1 for endolysosomal degradation enhanced the parasite’s ability to acquire the receptor and proliferate. TfR1 acquisition by *T. gondii* seems to have additional purpose for the parasite because the receptor is incorporated into newly forming daughter cells. Surprisingly, iron levels were not affected by the lack of functional TfR1 further supporting the idea that there is an alternative and essential function for the theft of these host proteins.

Iron acquisition by various pathogens occurs by direct iron scavenging or by breakdown of host iron homeostatic machinery ^10, 57, 58^. Direct iron scavenging is a method employed primarily by extracellular pathogens. Labile iron is traversed across the membranes of the pathogen via transporters such as Fec/Feo proteins in *Enterobacteria* or Yfe proteins in *Yersinia* ^41, 59, 60^. Several *T. gondii* genes were identified in this study and others (TGME49_267270, TGME49_266800, TGME49_261720, and TGME49_225530) that could function as such transporters. These genes would require functionality at the parasite PM, PVM, and potentially host PM interfaces for iron acquisition. Alternatively, iron may first be sequestered by siderophores ^10^. Generally, iron-laden siderophores are translocated into the pathogen, and iron is only then accessed after breakdown of the siderophore. This process requires siderophore synthesis, secretion, reuptake, breakdown, and regulation. In this study, we identified *T. gondii* genes with homology to siderophore iron trafficking, albeit from pathways of disparate siderophores (*P. aeruginosa* pyochelin, *Y. enterocolitica* yersiniabactin, and *Enterobacteria* enterobactin) ^61–63^.

Another prototypical mechanism of iron acquisition by pathogens is by breakdown of host iron-containing proteins ^58^. Heme is a primary target for such actions, hallmarked by pathogen possession of catabolism genes such as HemO in *Leptospira* ^64^. Other common targets are transferrin and lactoferrin. To access this source of iron, pathogens encode functional receptor homologs such as Transferrin-Binding Proteins and Lactoferrin Binding Proteins in *Trypanosoma, Neisseria, Haemophilus, and Bordatella* ^11–13, 65, 66^. These proteins compete for binding with transferrin and lactoferrin for sequestration and degradation to release iron for the pathogen. Yet another example is Etf3 in *Ehrlichia*, which is a ferritin-binding protein that leads to ferritinophagy to allow access of iron for the pathogen ^67^. In our study, we report no such homologs to any of these proteins encoded by *T. gondii*, yet, there was continued presence of the host iron-containing protein in the parasite.

Manipulation of host protein is a defining characteristic or viral and parasitic infections, but host protein ingestion, retention, and donation to offspring is a novel concept. New advances in proteomics reveal that virions such as those of influenza virus contain wide arrays of peptides derived from the host ^68^. Once thought to be artifactual incorporation during the viral lytic cycle, growing evidence suggests that the theft of host protein may be actively functional for pathogen-specific functions such as immune evasion ^69^.

In the apicomplexan parasite *Plasmodium*, reliance on heme scavenging for iron can be complemented by transferrin ^70^. The discovery that *Plasmodium* extracts contained a transferrin-binding protein indicated a potential parasite-encoded transferrin receptor ^49^. This *Plasmodium* TfR1 was shown to exhibit acylation distinct from human TfR1 ^71^ and the existence of this protein was debated ^72^. Here, we were unable to find a TfR1 homologous sequence in the *Plasmodium* genome. These observations suggest that *Plasmodium* also sequesters host transferrin receptor and this process is conserved in Apicomplexans.

The *T. gondii* PV membrane (PVM) is initially derived from the host PM lipids, but the host cell transmembrane proteins are excluded during invasion. This process is facilitated by extensive subversion of host CIN85, CD2AP, and ESCRT-I proteins ^4, 73^. After formation of the PV during *T. gondii* entry into the host cell, host-derived vesicles are detected within the PV lumen, and though their cargo remains ambiguous, TfR1 has been preferentially detected at the PVM interface ^74–76^. Moreover, *T. gondii* is capable of ingesting host cytosolic proteins *via* hijack of host ESCRT-II proteins ^6, 77^. Entry of host cell cytosolic cargo into tachyzoites is most likely mediated by a recently identified endocytosis function of the tachyzoite micropore ^7, 8^. It is not known whether entry of host TfR1, Tf, FHC, FLC, and CTLC proteins into tachyzoites is mediated by micropore function or mediated via a novel mechanism.

Theft of TfR1 and Tf by *T. gondii* presumptively requires vesicular trafficking from the surface of the host PM to the parasite. Recently, it has been discovered that the PV is selectively permissive to host vesicles by intravacuolar PVM budding that is Rab11a-dependent ^74^. In uninfected mammalian cells, after clathrin mediated endocytosis the Tf-TfR1 vesicle is acidified and either degraded by the endolysosome or recycled in the ERC as dictated by Rab11a and Rab11b ^29, 54, 78–81^. In uncommon events, we observed foci of TfR1 within the PV lumen that bore similarity to Rab11a foci, suggesting that TfR1 theft may occur through Rab11a modulation. The sialylation state of Tf and TfR1 may also determine the fate of Tf/TfR1 in this pathway ^82^, and the size of parasite-associated TfR1 exhibited signs of post-translational modifications in Western Blots. *T. gondii* also ingests host-CTLC which might indicate that rerouting of host endosomes occurs prior to acidification or recycling. However, enhancing TfR1 trafficking to the endolysosome with A24 resulted in increased parasite replication and TfR1 ingestion. Taken together these results may indicate that host Tf-TfR1 theft occurs prior to recycling. The parasite effector(s) mediating the hijack of this pathway and dictating the specificity of the cargo to be stolen is a high research priority for therapeutic intervention.

Tf-TfR1 theft may or may not be means of iron acquisition for *T. gondii.* Multiple methods to limit access of iron through genetic or protein manipulation of TfR1 could not completely prevent parasite replication. It is possible that there are redundant avenues for iron acquisition because there was also theft of host ferritin, or that the generational retention of TfR1 is long lived and resists short-term prevention of Tf-TfR1 theft. The iron content in parasites derived from TfR1 deficient cells was not reduced, but the reduced growth rate of the parasite within these cells could compensate and allow adequate levels of iron on a per parasite basis. While Tf-TfR1 theft has the potential to deliver iron, it is dispensable for this function, suggesting there is another novel and required function for parasite sequestration of host TfR1.

It is possible that theft of host TfR1 represents bulk theft of host cell membrane cargo. *T. gondii* has a high demand for membrane to build new PM and IMC membranes in daughter parasites during endodyogeny. These functions are involved in a fundamental aspect of *T. gondii* biology as TfR1 theft was observed in multiple virulence types, in multiple host species, and in developing endodyogenic daughter cells. Since the appearance of TfR1 in each of these cases is evocative of IMC morphology, it is also plausible that host proteins transiently function as structural component of the IMC in rapidly replicating tachyzoites. Bradyzoites which replicate rarely, or slowly, did not significantly accumulate host TfR1 indicating the cyst wall is a barrier to sequestering this host cell cargo.

Conversion to the bradyzoite life stage in immune-privileged sites is a means immune evasion that is not afforded to tachyzoites. Tachyzoites possess means of immune evasion as conversion-deficient strains are still not cleared and cause mortality ^39, 83^. Self- and non-self-recognition is a strong checkpoint in immunity that prevents attack of host cells. The accumulation and persistence of TfR1 in tachyzoites, but not bradyzoites, might correspond to the different needs of the two life stages. Recently, it was shown that *T. gondii* binds host PM protein CD36, and the host CD36 protein was required for parasite attachment and invasion of macrophages. Moreover, in the absence of host CD36 there was increased mortality of mice due to extensive tissue damage ^84^. Therefore, it is plausible that theft of TfR1, Tf, FHC/FLC, or tachyzoite binding of other host proteins such as CD36, may provide a cloak of host antigen associated with the tachyzoite PM as a mechanism to circumvent non-self recognition to prevent killing of host cells. In this possibility, the mechanism dictating the eventual loss of, or lack of maintaining, TfR1 during life stage conversion is a therapeutic target.

Overall, this study reports *Toxoplasma gondii* theft and incorporation of host proteins transferrin, transferrin receptor 1, ferritin heavy and light chains, and clathrin light chain into the tachyzoite PM or other compartments. This discovery reveals a novel host-pathogen interaction and supports emerging evidence that host proteins are repurposed for new functions in parasite biology.

## MATERIALS AND METHODS

### Tissue culture

MRC5 Human Lung Fibroblasts (ATCC: CCL-171) were maintained in Complete Media (Dulbecco’s Modified Eagles Medium with 4.5g/L glucose, L-glutamine, and sodium pyruvate, supplemented with 1X Penicillin/Streptomycin, Amphotericin B, and 10% Fetal Bovine Serum) at 37°C and 5% CO_2_. Cells were expanded when 85% confluent by detaching the cells with 0.05% Trypsin-EDTA and expanding cells at a ratio of 1:4 in terms of surface area available to the monolayer.

Chinese Hamster Ovary (CHO, ATCC: CCL-61) and CHO-derived cell lines were maintained in CHO Media (Ham’s F-12 Media supplemented with 1X Penicillin/Streptomycin, 2mg/mL D-Glucose, and 3% FBS) at 37°C and 5% CO_2_. CHO-derived TRVb cells were kindly gifted by Caroline A. Enns, Oregon Health & Science University. TRVb-derived hTfR, mTfR, and mTfR-R654A complemented cells, kindly gifted by Paul J. Schmidt, Boston Children’s Hospital and Harvard School of Medicine, were supplemented further with 1X Geneticin. Cells were expanded when 85% confluent by detaching the cells with 0.05% Trypsin-EDTA and expanding the cells at a ratio of 1:3.

Murine Fibroblasts (MFs) were isolated from C57BL/6J mice (Jackson Laboratories: 000664) adapted from established protocols ^85^. Briefly, 1cm-diameter sections from both ears of humanely euthanized mice were sterilized by submersion in 70% ethanol for 5 minutes and air dried in a sterile petri dish. The sections were digested at 37°C for 90 minutes in 2mL Digestion Buffer (2.5mg Pronase and 5mg Collagenase D in Isocove’s Complete Media). Cells were released from tissue by straining through a 70µm cell strainer and washed in 10mL MF Media (Isocove’s DMEM with L-Glutamine and 25mM HEPES supplemented with 1X Penicillin/ Streptomycin, Amphotericin B, Non-Essential Amino Acids, 0.1mM Sodium Pyruvate, 0.2mM Glutamine XL, 10% Fetal Bovine Serum, and 0.1mM 2-mercaptoethanol) before seeding into tissue-culture treated plates. Cells were regularly maintained in MF Media at 37°C and 5% CO_2_ and expanded when 80% confluent by detaching the cells with 0.05% Trypsin with EDTA, expanding cells at a ratio of 1:4 in terms of surface area available to the monolayer.

Type I Strain RH (ATCC: 50174) and Type II Strain Prugniaud (a kind gift from Dr. David J. Bzik, Dartmouth) *Toxoplasma gondii* parasites were regularly maintained by regular passage upon lysis in MRC5 fibroblasts at an MOI of ∼0.5 in Black Media (Dulbecco’s Modified Eagles Medium with 4.5g/L glucose, L-glutamine, and sodium pyruvate, supplemented with 1X Penicillin/Streptomycin, Amphotericin B, and 1% Fetal Bovine Serum) at 37°C and 5% CO2. 10,000 Prugniaud strain tachyzoites were infected intraperitoneally into C57BL/6J mice every 10-12 *in vitro* passages, and re-isolated from the mouse brains to ensure maintenance of the cyst-forming ability of this strain of parasite. Type II Strain ME49 (ATCC: 50611) was routinely passaged by cyst isolation from 5 week-post-infected CBA/J mice brains and subsequent inoculation of 10 cysts intragastrointestinally.

All tissue culture experiments were performed when fibroblasts reached 100% confluency. Where appropriate, (Z)-4-hydroxy Tamoxifen (Cayman Chemical Company #14854) was dissolved in DMSO at 2.5mM and added to cultures to a final concentration of 2.5µM at the indicated timepoints. For A24 monoclonal antibody blockade experiments, A24 (Creative Biolabs #TAB-622CT) was administered to cultures at 1µg/mL every two days starting 15 minutes before infection.

### Animal Maintenance

All animal husbandry was performed by the laboratory of Jason P. Gigley at the University of Wyoming and in accordance with the guidelines approved by the University of Wyoming Institutional Animal Care and Use Committee. C57BL/6J (#000664) and Ubiquitin-CreER^T2^ (#007001) mice were purchased from Jackson Laboratories. TfR1^Flx^ mice were a kind gift from Mark Andrews, Harvard University. Mouse genotypes were assessed from genomic DNA isolated from ear clips *via* digestion in DirectPCR Digestion Buffer + Proteinase K (Viagen Biotech) using the following primer pairs: Ubiquitin-CreER^T2^ Forward (0.5µM PCR Reaction): 5’-GAC GTC ACC CGT TCT GTT G -3’, Ubiquitin-CreER^T2^ Reverse (0.5µM PCR Reaction): 5’-AGG CAA ATT TTG GTG TAC GG -3’, and TfR^Flx^ Forward (0.3µM PCR Reaction): (5’-TTC AGT TCC CAG TGA CCA CA -3’), TfR^Flx^ Reverse (0.3µM PCR Reaction): 5’-TCC TTT CTG TGC CCA GTT CT -3’, where the presence of the Ub-CreER^T2^ transgene was indicated by a 475nt agarose gel band, the wild-type TfR1 gene was indicated by a 165nt agarose gel band, and the floxed TfR1 gene was indicated by a 310nt agarose gel band.

### Cyst Isolation and Reactivation

Cyst isolation was adapted from a protocol described previously ^86^. Chronically infected mice were humanely euthanized by CO2 inhalation and dissected with sterile instruments to remove the brain tissue, which was subsequently homogenized with a Dounce tissue grinder in PBS. Cysts were manually enumerated by sampling 10µL aliquots of brain homogenate under a light microscope. Cysts were further purified on a 90/40/20% Percoll gradient in 1X PBS and retrieved from the 90/40 interface following differential centrifugation at 900 RCF for 20 minutes at 21°C with no brake. Bradyzoites were released by suspension in 100µL 0.05% Trypsin-EDTA for 30 seconds, quenched in Black Media, and immediately inoculated into *in vitro* cultures or subjected to flow cytometry protocol. For microscopic evaluation, reactivated parasites were subjected to confocal microscopy methodology 24-36 hours after *in vitro* inoculation.

### Brain Sectioning

Brain sectioning protocol was adapted from methods described previously ^87^. C57BL/6 mice chronically infected with ME49 strain *T. gondii* were anesthetized with a barbiturate cocktail and exsanguinated *via* perfusion. Perfusion was performed by a peristaltic pump; the 27G needle outlet was inserted into the left ventricle of the heart, an incision was made through the right atrium, and the pump operated at a rate of 10mL / minute. Perfusion occurred first with 20mL of saline (25U Heparin / mL, 0.9% (w/v NaCl), then with 100mL of fixative (4% w/v Paraformaldehyde in 0.1M phosphate buffer). The mouse corpse was incubated at 4°C for 24h in fixative, and the whole brain incubated a further five days in cryopreservative (10% v/v glycerol, 2.5% v/v dimethyl sulfoxide in 0.1M phosphate buffer). The brains were mounted in OCT solution and affixed to a microtome by freezing with dry ice. 40µm sections of brain were transversely collected and stored at 4°C in 0.1M phosphate buffer until prepared for microscopic evaluation.

### Confocal Microscopy (CM)

Coverslips (#1 Thickness German Glass, 15mm Diameter) were individually washed in a 1N HCl solution for 24 hours. The coverslips were individually and subsequently rinsed in 70%, 80%, and 90% ethanol solutions before storage in 95% ethanol. Fibroblasts were inoculated onto prepared and dried coverslips upon expansion into a 24-well format. Immediately upon reaching confluency on the coverslips, fibroblasts were inoculated with 1,000 freshly lysed parasites per coverslip for 24 hours.

Infected coverslips were fixed, permeabilized and blocked, and stained for immunofluorescence (IFA) in the culture wells before mounting on a microscope slide. Fixation occurred in 4% Paraformaldehyde (w/v) in PHEM buffer (60mM PIPES, 27mM HEPES, 10mM EGTA, and 8.25mM MgSO4) for 10 minutes at ambient temperature. The fixative was washed away and residual fixative was quenched in 0.1M glycine. The cells were permeabilized and blocked in PHEMBlock (3% FBS and 0.2% Triton X-100 in PHEM buffer) for 30 minutes at room temperature. Host proteins were first stained by incubation with indicated rabbit IgG in PHEMBlock for 16 hours at 4°C in the dark, then counterstained with goat IgG secondary antibody in PHEMBlock for 8 hours at 4°C in the dark. Parasite proteins or alternative host proteins were stained next by incubation with indicated IgG in PHEMBlock for 16 hours at 4°C, followed by counterstain with indicted IgG secondary antibody or streptavidin-conjugated fluorophores in PHEMBlock for 8 hours at 4°C in the dark. After staining was complete, coverslips were washed once in 2µM DAPI in PBS and twice in PBS. Coverslips were mounted on microscope slides with Prolong Gold Antifade Mountant with DAPI (ThermoFisher P36931).

For alternative permeabilization experiments, coverslips were prepared, seeded, inoculated, fixed, and quenched as above. Permeabilization and blocking occurred by a 30 minute incubation in 0.2% (v/v) Triton X-100, 0.01% (w/v) Saponin, or 0.005% (w/v) in 3% FBS PHEM. All staining steps occurred identically to above except the antibodies were diluted in 3% FBS PHEM without permeabilization agent.

Fixed brain sections were blocked in blocking buffer (3% FBS and 0.1% Triton X-100 in PBS) for one hour and stained with concurrent DBA-FITC and α-TfR1 (JF0956) in blocking buffer overnight at 4°C. Sections were washed three times with PBS and secondarily stained with goat α-rabbit secondary antibody in blocking buffer for eight hours at 4°C. Brains were washed with 2µg/mL DAPI in PBS once, and twice in PBS before mounting on a microscope slide with Prolong Glass Antifade Mountant (ThermoFisher P36984).

All samples were imaged with an Olympus IX81 confocal microscope on Metamorph v7.8.6.0 software and analyzed with ImageJ software v1.53f51. Images may have modified brightness and contrast for presentation purposes.

### Real-Time PCR (RT-PCR)

Fibroblast lines were inoculated with *T. gondii* according to host cell type; MRC5, MF, or CHO lines were infected with 100 tachyzoites for six days in Black Media. Host cells and parasites were lifted with a cell scraper and pelleted by centrifugation. Genomic DNA was purified from the samples using the DNeasy Blood & Tissue Kit (Qiagen 69506) and normalized to 6.25ng/µL. A standard curve of known quantity of RH gDNA was assessed alongside 50ng of sample DNA as template to query for the *T. gondii* unique gene B1 using SSO Advanced SYBR Master Mix (Bio-Rad 1725017) and 0.5µM primers (F: 5’- CGT CCG TCG TAA TAT CAG -3’, R: 5’- GAC TTC ATG GGA CGA TAT G -3’). The thermocycler conditions were 1: 95°C for 10 minutes, 2: 95°C for 5 seconds, 3: 60°C for 5 seconds, 4: 72°C for 5 seconds + plate read, 5: Cycle 2-4, 44 repetitions, 6: Melt Curve 65°C to 95°C with 0.5°C, 5 second increments + plate read.

### Quantitative Reverse Transcriptase PCR (qRT-PCR)

Samples were dissolved in TRI reagent (Sigma-Aldrich T9424), further disrupted in a tissue grinder, and phase-separated by addition of chloroform in a 1:5 ratio. The aqueous RNA was precipitated in isopropanol and washed in 75% ethanol, air-dried, and resuspended in DEPC-Treated water supplemented with 1X RNaseOUT (Invitrogen 10777019). RNA samples were further purified by the RNA Cleanup protocol of RNeasy Kit (Qiagen 74104). 1µg of RNA samples were used as template in cDNA generation using Oligo d(t) primers and Superscript IV (Invitrogen 18091050) according to manufacturer instructions. PCR of the resulting cDNA was performed with SSO Advanced SYBR Master Mix (Bio-Rad 1725017) and 0.5µM of primers against mouse actin (F: 5’- ACC CAC ACT GTG CCC TAC TA -3’, R: 5’- CAC GCT CGG TCA GGA TCT TC -3’) ^88^ and mouse TfR1 (F: 5’- TCA AGC CAG ATC AAT TCT C -3’, R: 5’- AGC CAG TTT CAT CTC CAC ATG -3’) ^89^. The thermocycler conditions were 1: 95°C for 3 minutes, 2: 95°C for 15 seconds, 3: 60°C for 15 seconds, 4: 72°C for 1 minute 20 seconds + plate read, 5: Cycle 2-4, 44 repetitions, 6: Melt Curve 65°C to 95°C with 0.5°C, 5 second increments + plate read. Data were analyzed by the ΔΔCt method.

### Western Blot (WB)

Fibroblast lines or lines infected with *T. gondii* at an MOI of 0.1 were incubated in Black Media for two days at 37°C and 5% CO2. Cells were detached from the flask with a sterile cell scraper. Only for infected flasks, parasites were isolated by disrupting host cells by twice-passage of samples through a 27G needle and twice-filtration through a 3µm membrane. Every sample was washed twice by centrifugation in PBS, and finally resuspended in Lysis Buffer (20mM HEPES, 10mM Sodium Citrate, 10mM Sodium Pyrophosphate, 50mM Sodium Fluoride, 1mM Sodium Vanadate, 50mM B-Glycerophosphate, 5mM EDTA, and 0.5% NP-40 + fresh addition of 2mM Benzamidine, 0.2mM PMSF, 40µg/mL Leupeptin, 40µg/mL Aprotinin, and 4µg/mL Pepstatin A). Cell lysate was agitated with a tissue grinder and insoluble material was removed by centrifugation. 30µg of purified protein was denatured in SDS Loading Buffer (4% w/v SDS, 20% v/v Glycerol, 0.1M Tris-HCl pH 6.8, 2mg/mL bromophenol blue, and 0.2M DTT) for 10 minutes at 95°C before loading in a 10% SDS-PAGE Gel (Bio-Rad 4561034). Sample proteins were separated in 1X SDS Running Buffer (25mM Tris, 20mM Glycine, 0.1% w/v SDS) at 13mA for ∼1.5hr until the reference ladder (Bio-Rad 161-0374) was separated. Transfer was completed to a methanol-activated PVDF membrane in Transfer Buffer (20% v/v methanol in 25mM Tris, 192mM glycine) at 300mA and 4°C for 2hr. Membranes were blocked in 5% (w/v) Nonfat Milk in Wash Buffer (0.1% Tween-20 in PBS) for 30 minutes, washed three times, blotted with primary antibody in Blocking Buffer for 1.5hr at 22°C, washed three times, blotted with secondary antibody in Blocking Buffer for 1.5hr at 22°C, and washed 4 times. Immediately before chemiluminescent exposure in an Invitrogen iBright machine, membranes were developed in a solution of equal volumes of ECL peroxidase and luminol reagents (Bio-Rad 170-5060).

### Inductively Coupled Plasma Mass Spectometry (ICPMS)

10^5^ RH strain *T. gondii* tachyzoites were inoculated into fibroblasts and grown at 37°C and 5% CO2. Prior to lysis, monolayers were lifted with a cell scraper, passaged twice through a 27G needle and twice through a 3µm filter to isolate parasites. Unless otherwise stated, 5*10^6^ isolated parasites were digested in 100µL concentrated HNO3 for 45 minutes at 94°C and diluted with 900µL 1% (v/v) HNO3 in dH2O before ICPMS. Analysis was performed using an Agilent 8900 triple quad equipped with an SPS4 autosampler. The system was operated at a radio frequency power of 1550 W, an argon plasma gas flow rate of 15 L/min, and Ar carrier gas flow rate of 0.9 L/min. Data were quantified using weighed, serial dilutions of a multi-element standard (CEM 2, (VHG labs, VHG-SM70B-100) for Mg, Mn, Fe, Cu, and Zn, and single element stands for S (SpexCertiPrep, PLP9-2M) and P (SpexCertiPrep, PLP9-2Y). Data were quantified using a 12-point calibration curve.

### Flow Cytometry

Tachyzoites were inoculated into indicated fibroblast culture and later retrieved by mechanical disruption and filtration, similar as in Western Blot protocol. Alternatively, bradyzoites isolated from cyst isolation and reactivation protocol were retrieved after the trypsinization quench. Parasites were enumerated on a hemacytometer and plated at 10^6^ per well in a FACS plate. Viability was assessed by Live/Dead Aqua (Invitrogen L34966), according to manufacturer instructions, prior to fixation in 4% paraformaldehyde in PHEM. There were four separate rounds of antibody staining containing, in order, α-TfR1 JF0956, Goat α-Rabbit IgG Secondary, α-Gra5, and Donkey α-Mouse IgG Secondary in PHEMBlock. Cells were finally resuspended in PBS and processed in Guava EasyCyte 12HT Flow Cytometer Serial #8470120112 and analyzed in FlowJo v.9.7.7 software.

### In silico identification of potential T. gondii iron-related genes

Iron-related proteins were manually selected and cataloged for query by literature review. FASTA protein sequences of reference genes identified from primary reports were downloaded from the NCBI Protein Database and queried for homology in the *Toxoplasma gondii* ME49 genome (ToxoDB Toxoplasma Bioinformatics Resource, ToxoDB.org) or the *Plasmodium falciparum* 3D7 genome (PlasmoDB Plasmodium Bioinformatics Resource, PlasmoDB.org) by pBLAST. Potential candidates were limited to those that had bit scores greater than 50 and E-values less than 1.00*10^-5^. Identified *T. gondii* genes were then cross-referenced to CRISPR Fitness Scoring ^33^. Any protein sequence alignments were performed by ClustalW analysis with a gap penalty of 10.0 in MacVector v17.0.10 software.

### Antibodies and Reagents Used

α-TfR1 (Clone JF0956) Rabbit IgG, Invitrogen MA5-32500 (Diluted 1:1000 CM, 1:500 FC, 1:1000 Brain Sections).

α-Sag1 (Clone D16S) Mouse IgG, Invitrogen MA5-18628 (Diluted 1:500 CM, 1:200 WB, 1:500 FC)

α-Gra5 Mouse IgG, Biotem TG 17-113 (Diluted 1:500 CM, 1:250 FC)

α-Tf (Clone F2G8H6) Mouse IgG, Invitrogen MA5-14690 (Diluted 1:250 CM)

Biotin-α-Tf (Polyclonal) Goat IgG, Invitrogen Bethyl Laboratories A80-128B (Diluted 1:250 CM)

α-TfR1 (Clone H68.4) Mouse IgG, Invitrogen 13-6800 (Diluted 1:500 CM, WB)

α-CTLC (Clone 232-340) Mouse IgG, Abnova H00001213-M05 (Diluted 1:250 CM)

α-GAPDH (Polyclonal) Rabbit IgG, Proteintech 10494-1-AP (Diluted 1:2000 WB)

Goat α-Rabbit IgG (H+L) AlexaFluor594, Invitrogen A11012 (Same dilution as primary) Goat α-Rabbit IgG (H+L) AlexaFluor405, Invitrogen A31556 (Same dilution as primary)

Goat α-Rabbit IgG (H+L), HRP Conjugate, Invitrogen G21234 (1mg/mL in PBS Stock, 1:2000 WB) Donkey α-Mouse IgG (H+L) AlexaFluor 488, Invitrogen A21202 (Same dilution as primary) Donkey α-Mouse IgG (H+L), HRP Conjugate, Invitrogen SA1-100 (1:2000 WB)

Streptavidin-AlexaFluor594 Conjugate, Invitrogen S11227 (1mg/mL in 5mM NaN3 in PBS Stock, same dilution as biotinylated target)

Dolichos Bifluorus Agglutinin, FITC Conjugate, Invitrogen L32474 (2µg/mL CM)

### Statistics

Statistics and graph generation were completed using Graphpad Prism v.9 and Microsoft Excel v16.54 software. Statistics were performed between two groups by Student’s t-test, between multiple groups by Ordinary One-Way ANOVA, or between two groups within multiple conditions by Two-Way ANOVA. Relevant statistical values such as sample size (n), P values (p), t-values (t), F-values (F), and degrees of freedom (df) are provided in figures and figure legends. Figures were assembled in Adobe Illustrator 2021, references in Endnote X8.

### Data Availability

The authors declare that all data supporting the findings of this study are available within the paper and its supplementary information files.

## Supporting information

References for suplemental table 1

z stack

z stack

z stack

## Acknowledgements

We thank Caroline Enns, Paul Schmidt, Mark Andrews, David Bzik, Barbara Fox, Sarah Ewald, Daniel Wall, Donald Jarvis, Jay Gatlin, and Jonathan Fox for donation of reagents and equipment use. Funding was provided by UW INBRE Pilot Grant awarded to J.P.G. and UW INBRE Graduate Assistantship awarded to S.L.D. under NIH Grant # 2P20GM103432. Additional funding was provided by NIH Grants #R21AI1592 and #R21AI1612 awarded to J.P.G. ICPMS measurements were performed in the OHSU Elemental Analysis Core with partial support from NIH (S10OD028492).

## Author Contributions

The principal investigator and corresponding author of this project is J.P.G. Project conception was conducted by S.L.D. and J.P.G. Experiments were designed by S.L.D. and performed by S.L.D., A.M., and L.L.N. Data analysis was conducted by S.L.D. Intellectual contributions were made by S.L.D., J.P.G., B.A.F., D.J.B., and M.P.S. Technical expertise was provided by J.P.G., D.J.B., and M.P.S. Manuscript preparation was performed by S.L.D. and J.P.G., with contributions by all authors on revisions. Project management was performed by J.P.G.

## Materials *and Correspondence*

Material requests or other correspondence should be addressed to Jason P. Gigley at or 1000 University Ave, University of Wyoming, 6005 AgC, Laramie, Wyoming, USA.

## REFERENCES

1. Phillips MA, Burrows JN, Manyando C, van Huijsduijnen RH, Van Voorhis WC, Wells TNC. Malaria. Nature Reviews Disease Primers 3, 17050 (2017).

2. Gibson AR, Striepen B. Cryptosporidium. Current biology : CB 28, R193–r194 (2018).

3. Dubey JP, Lindsay DS, Speer CA. Structures of Toxoplasma gondii tachyzoites, bradyzoites, and sporozoites and biology and development of tissue cysts. Clinical microbiology reviews 11, 267–299 (1998).

4. Schwab JC, Beckers CJ, Joiner KA. The parasitophorous vacuole membrane surrounding intracellular Toxoplasma gondii functions as a molecular sieve. Proceedings of the National Academy of Sciences of the United States of America 91, 509–513 (1994).

5. Coppens I, et al. Toxoplasma gondii Sequesters Lysosomes from Mammalian Hosts in the Vacuolar Space. Cell 125, 261–274 (2006).

6. Rivera-Cuevas Y, et al. Toxoplasma gondii exploits the host ESCRT machinery for parasite uptake of host cytosolic proteins. PLoS pathogens 17, e1010138 (2021).

7. Wan W, et al. The Toxoplasma micropore mediates endocytosis for selective nutrient salvage from host cell compartments. Nat Commun 14, 977 (2023).

8. Koreny L, et al. Stable endocytic structures navigate the complex pellicle of apicomplexan parasites. Nat Commun 14, 2167 (2023).

9. Lane DJ, et al. Cellular iron uptake, trafficking and metabolism: Key molecules and mechanisms and their roles in disease. Biochim Biophys Acta 1853, 1130–1144 (2015).

10. Miethke M, Marahiel MA. Siderophore-based iron acquisition and pathogen control. Microbiology and molecular biology reviews : MMBR 71, 413–451 (2007).

11. Schryvers AB, Morris LJ. Identification and characterization of the transferrin receptor from Neisseria meningitidis. Molecular microbiology 2, 281–288 (1988).

12. Menozzi FD, Gantiez C, Locht C. Identification and purification of transferrin- and lactoferrin-binding proteins of Bordetella pertussis and Bordetella bronchiseptica. Infection and immunity 59, 3982–3988 (1991).

13. Kabiri M, Steverding D. Trypanosoma brucei transferrin receptor: Functional replacement of the GPI anchor with a transmembrane domain. Molecular and biochemical parasitology 242, 111361 (2021).

14. Clark MA, et al. Host iron status and iron supplementation mediate susceptibility to erythrocytic stage Plasmodium falciparum. 5, 4446 (2014).

15. Oliveira MC, Coutinho LB, Almeida MPO, Briceno MP, Araujo ECB, Silva NM. The Availability of Iron Is Involved in the Murine Experimental Toxoplasma gondii Infection Outcome. Microorganisms 8, (2020).

16. Fox BA, Gigley JP, Bzik DJ. Toxoplasma gondii lacks the enzymes required for de novo arginine biosynthesis and arginine starvation triggers cyst formation. International journal for parasitology 34, 323–331 (2004).

17. Krug EC, Marr JJ, Berens RL. Purine metabolism in Toxoplasma gondii. J Biol Chem 264, 10601–10607 (1989).

18. Romano JD, Sonda S, Bergbower E, Smith ME, Coppens I. Toxoplasma gondii salvages sphingolipids from the host Golgi through the rerouting of selected Rab vesicles to the parasitophorous vacuole. Mol Biol Cell 24, 1974–1995 (2013).

19. Aghabi D, Sloan M, Dou Z, Guerra AJ, Harding CR. The vacuolar iron transporter mediates iron detoxification in Toxoplasma gondii. bioRxiv, (2021).

20. Dimier IH, Bout DT. Interferon-gamma-activated primary enterocytes inhibit Toxoplasma gondii replication: a role for intracellular iron. Immunology 94, 488–495 (1998).

21. Pamukcu S, Cerutti A, Bordat Y, Hem S, Rofidal V, Besteiro S. Differential contribution of two organelles of endosymbiotic origin to iron-sulfur cluster synthesis and overall fitness in Toxoplasma. PLoS pathogens 17, e1010096 (2021).

22. Bergmann A, et al. Toxoplasma gondii requires its plant-like heme biosynthesis pathway for infection. PLoS pathogens 16, e1008499 (2020).

23. Zhang X, et al. Functional characterization of a unique cytochrome P450 in Toxoplasma gondii. Oncotarget 8, 115079–115088 (2017).

24. Sibley LD, Lawson R, Weidner E. Superoxide dismutase and catalase in Toxoplasma gondii. Molecular and biochemical parasitology 19, 83–87 (1986).

25. Brydges SD, Carruthers VB. Mutation of an unusual mitochondrial targeting sequence of SODB2 produces multiple targeting fates in Toxoplasma gondii. J Cell Sci 116, 4675–4685 (2003).

26. Aw YTV, et al. A key cytosolic iron-sulfur cluster synthesis protein localizes to the mitochondrion of Toxoplasma gondii. Molecular microbiology 115, 968–985 (2021).

27. Seidi A, et al. Elucidating the mitochondrial proteome of Toxoplasma gondii reveals the presence of a divergent cytochrome c oxidase. Elife 7, (2018).

28. Hanover JA, Beguinot L, Willingham MC, Pastan IH. Transit of receptors for epidermal growth factor and transferrin through clathrin-coated pits. Analysis of the kinetics of receptor entry. J Biol Chem 260, 15938–15945 (1985).

29. Giannetti AM, Halbrooks PJ, Mason AB, Vogt TM, Enns CA, Bjorkman PJ. The molecular mechanism for receptor-stimulated iron release from the plasma iron transport protein transferrin. Structure 13, 1613–1623 (2005).

30. Kolachala VL, Sesikeran B, Nair KM. Evidence for a sequential transfer of iron amongst ferritin, transferrin and transferrin receptor during duodenal absorption of iron in rat and human. World J Gastroenterol 13, 1042–1052 (2007).

31. Renaud EA, et al. Disrupting the plastidic iron-sulfur cluster biogenesis pathway in *Toxoplasma gondii* has pleiotropic effects irreversibly impacting parasite viability. J Biol Chem 298, 102243 (2022).

32. Almeida MPO, Ferro EAV, Briceño MPP, Oliveira MC, Barbosa BF, Silva NM. Susceptibility of human villous (BeWo) and extravillous (HTR-8/SVneo) trophoblast cells to Toxoplasma gondii infection is modulated by intracellular iron availability. Parasitol Res 118, 1559–1572 (2019).

33. Sidik SM, et al. A Genome-wide CRISPR Screen in Toxoplasma Identifies Essential Apicomplexan Genes. Cell 166, 1423–1435 e1412 (2016).

34. Lill R. Function and biogenesis of iron-sulphur proteins. Nature 460, 831–838 (2009).

35. Scharton-Kersten TM, Yap G, Magram J, Sher A. Inducible nitric oxide is essential for host control of persistent but not acute infection with the intracellular pathogen Toxoplasma gondii. The Journal of experimental medicine 185, 1261–1273 (1997).

36. Khan IA, Schwartzman JD, Matsuura T, Kasper LH. A dichotomous role for nitric oxide during acute Toxoplasma gondii infection in mice. Proceedings of the National Academy of Sciences of the United States of America 94, 13955–13960 (1997).

37. Jun CD, Kim SH, Soh CT, Kang SS, Chung HT. Nitric oxide mediates the toxoplasmastatic activity of murine microglial cells in vitro. Immunological investigations 22, 487–501 (1993).

38. Chao CC, Anderson WR, Hu S, Gekker G, Martella A, Peterson PK. Activated microglia inhibit multiplication of Toxoplasma gondii via a nitric oxide mechanism. Clinical immunology and immunopathology 67, 178–183 (1993).

39. Waldman BS, Schwarz D, Wadsworth MH, 2nd, Saeij JP, Shalek AK, Lourido S. Identification of a Master Regulator of Differentiation in Toxoplasma. Cell 180, 359–372 e316 (2020).

40. Abboud S, Haile DJ. A novel mammalian iron-regulated protein involved in intracellular iron metabolism. J Biol Chem 275, 19906–19912 (2000).

41. Makui H, Roig E, Cole ST, Helmann JD, Gros P, Cellier MF. Identification of the Escherichia coli K-12 Nramp orthologue (MntH) as a selective divalent metal ion transporter. Molecular microbiology 35, 1065–1078 (2000).

42. Sahu T, et al. ZIPCO, a putative metal ion transporter, is crucial for Plasmodium liver-stage development. EMBO Mol Med 6, 1387–1397 (2014).

43. Shindo M, et al. Functional role of DMT1 in transferrin-independent iron uptake by human hepatocyte and hepatocellular carcinoma cell, HLF. Hepatol Res 35, 152–162 (2006).

44. Ford GC, et al. Ferritin: design and formation of an iron-storage molecule. Philosophical transactions of the Royal Society of London Series B, Biological sciences 304, 551–565 (1984).

45. Donovan A, et al. The iron exporter ferroportin/Slc40a1 is essential for iron homeostasis. Cell metabolism 1, 191–200 (2005).

46. Breinich MS, et al. A dynamin is required for the biogenesis of secretory organelles in Toxoplasma gondii. Current biology : CB 19, 277–286 (2009).

47. Fox BA, et al. Toxoplasma gondii Parasitophorous Vacuole Membrane-Associated Dense Granule Proteins Orchestrate Chronic Infection and GRA12 Underpins Resistance to Host Gamma Interferon. mBio 10, (2019).

48. Beckers CJ, Dubremetz JF, Mercereau-Puijalon O, Joiner KA. The Toxoplasma gondii rhoptry protein ROP 2 is inserted into the parasitophorous vacuole membrane, surrounding the intracellular parasite, and is exposed to the host cell cytoplasm. J Cell Biol 127, 947–961 (1994).

49. Rodriguez MH, Jungery M. A protein on Plasmodium falciparum-infected erythrocytes functions as a transferrin receptor. Nature 324, 388–391 (1986).

50. Sheffield HG, Melton ML. The fine structure and reproduction of Toxoplasma gondii. The Journal of parasitology 54, 209–226 (1968).

51. Nishi M, Hu K, Murray JM, Roos DS. Organellar dynamics during the cell cycle of Toxoplasma gondii. J Cell Sci 121, 1559–1568 (2008).

52. McGraw TE, Greenfield L, Maxfield FR. Functional expression of the human transferrin receptor cDNA in Chinese hamster ovary cells deficient in endogenous transferrin receptor. J Cell Biol 105, 207–214 (1987).

53. Schmidt PJ, Toran PT, Giannetti AM, Bjorkman PJ, Andrews NC. The transferrin receptor modulates Hfe-dependent regulation of hepcidin expression. Cell metabolism 7, 205–214 (2008).

54. Sharma M, Giridharan SS, Rahajeng J, Naslavsky N, Caplan S. MICAL-L1 links EHD1 to tubular recycling endosomes and regulates receptor recycling. Mol Biol Cell 20, 5181–5194 (2009).

55. Lepelletier Y, et al. Prevention of mantle lymphoma tumor establishment by routing transferrin receptor toward lysosomal compartments. Cancer Res 67, 1145–1154 (2007).

56. Skariah S, McIntyre MK, Mordue DG. Toxoplasma gondii: determinants of tachyzoite to bradyzoite conversion. Parasitol Res 107, 253–260 (2010).

57. Lau CK, Krewulak KD, Vogel HJ. Bacterial ferrous iron transport: the Feo system. FEMS Microbiol Rev 40, 273–298 (2016).

58. Choby JE, Skaar EP. Heme Synthesis and Acquisition in Bacterial Pathogens. J Mol Biol 428, 3408–3428 (2016).

59. Kammler M, Schon C, Hantke K. Characterization of the ferrous iron uptake system of Escherichia coli. Journal of bacteriology 175, 6212–6219 (1993).

60. Bearden SW, Perry RD. The Yfe system of Yersinia pestis transports iron and manganese and is required for full virulence of plague. Molecular microbiology 32, 403–414 (1999).

61. Serino L, Reimmann C, Visca P, Beyeler M, Chiesa VD, Haas D. Biosynthesis of pyochelin and dihydroaeruginoic acid requires the iron-regulated pchDCBA operon in Pseudomonas aeruginosa. Journal of bacteriology 179, 248–257 (1997).

62. Nahlik MS, Fleming TP, McIntosh MA. Cluster of genes controlling synthesis and activation of 2,3-dihydroxybenzoic acid in production of enterobactin in Escherichia coli. Journal of bacteriology 169, 4163–4170 (1987).

63. Fetherston JD, Bertolino VJ, Perry RD. YbtP and YbtQ: two ABC transporters required for iron uptake in Yersinia pestis. Molecular microbiology 32, 289–299 (1999).

64. Murray GL, Ellis KM, Lo M, Adler B. Leptospira interrogans requires a functional heme oxygenase to scavenge iron from hemoglobin. Microbes and infection 10, 791–797 (2008).

65. Schryvers AB. Identification of the transferrin- and lactoferrin-binding proteins in Haemophilus influenzae. Journal of medical microbiology 29, 121–130 (1989).

66. Schryvers AB, Morris LJ. Identification and characterization of the human lactoferrin-binding protein from Neisseria meningitidis. Infection and immunity 56, 1144–1149 (1988).

67. Yan Q, et al. Iron robbery by intracellular pathogen via bacterial effector-induced ferritinophagy. Proceedings of the National Academy of Sciences of the United States of America 118, (2021).

68. Shaw ML, Stone KL, Colangelo CM, Gulcicek EE, Palese P. Cellular proteins in influenza virus particles. PLoS pathogens 4, e1000085 (2008).

69. Berri F, et al. Annexin V incorporated into influenza virus particles inhibits gamma interferon signaling and promotes viral replication. J Virol 88, 11215–11228 (2014).

70. Pollack S, Fleming J. Plasmodium falciparum takes up iron from transferrin. Br J Haematol 58, 289–293 (1984).

71. Haldar K, Henderson CL, Cross GA. Identification of the parasite transferrin receptor of Plasmodium falciparum-infected erythrocytes and its acylation via 1,2-diacyl-sn-glycerol. Proceedings of the National Academy of Sciences 83, 8565 (1986).

72. Pollack S, Schnelle V. Inability to detect transferrin receptors on P. falciparum parasitized red cells. Br J Haematol 68, 125–129 (1988).

73. Guerin A, et al. Efficient invasion by Toxoplasma depends on the subversion of host protein networks. Nat Microbiol 2, 1358–1366 (2017).

74. Hartman EJ, Asady B, Romano JD, Coppens I. The Rab11-family interacting proteins reveal selective interaction of mammalian recycling endosomes with the Toxoplasma parasitophorous vacuole in a Rab11- and Arf6-dependent manner. Mol Biol Cell 33, ar34 (2022).

75. Romano JD, et al. The parasite Toxoplasma sequesters diverse Rab host vesicles within an intravacuolar network. J Cell Biol 216, 4235–4254 (2017).

76. Cygan AM, et al. Proximity-Labeling Reveals Novel Host and Parasite Proteins at the Toxoplasma Parasitophorous Vacuole Membrane. mBio 12, e0026021 (2021).

77. Dou Z, McGovern OL, Di Cristina M, Carruthers VB. Toxoplasma gondii ingests and digests host cytosolic proteins. mBio 5, e01188–01114 (2014).

78. Schafer JC, McRae RE, Manning EH, Lapierre LA, Goldenring JR. Rab11-FIP1A regulates early trafficking into the recycling endosomes. Exp Cell Res 340, 259–273 (2016).

79. Ullrich O, Reinsch S, Urbe S, Zerial M, Parton RG. Rab11 regulates recycling through the pericentriolar recycling endosome. J Cell Biol 135, 913–924 (1996).

80. Ren M, Xu G, Zeng J, De Lemos-Chiarandini C, Adesnik M, Sabatini DD. Hydrolysis of GTP on rab11 is required for the direct delivery of transferrin from the pericentriolar recycling compartment to the cell surface but not from sorting endosomes. Proceedings of the National Academy of Sciences of the United States of America 95, 6187–6192 (1998).

81. Schlierf B, Fey GH, Hauber J, Hocke GM, Rosorius O. Rab11b is essential for recycling of transferrin to the plasma membrane. Exp Cell Res 259, 257–265 (2000).

82. Snider MD, Rogers OC. Intracellular movement of cell surface receptors after endocytosis: resialylation of asialo-transferrin receptor in human erythroleukemia cells. J Cell Biol 100, 826–834 (1985).

83. Licon MH, et al. A positive feedback loop controls Toxoplasma chronic differentiation. Nat Microbiol 8, 889–904 (2023).

84. Zhao Y, Reyes J, Rovira-Diaz E, Fox BA, Bzik DJ, Yap GS. Cutting Edge: CD36 Mediates Phagocyte Tropism and Avirulence of Toxoplasma gondii. Journal of immunology (Baltimore, Md : 1950) 207, 1507–1512 (2021).

85. Khan M, Gasser S. Generating Primary Fibroblast Cultures from Mouse Ear and Tail Tissues. J Vis Exp, (2016).

86. Watts EA, Dhara A, Sinai AP. Purification Toxoplasma gondii Tissue Cysts Using Percoll Gradients. Curr Protoc Microbiol 45, 20C 22 21–20C 22 19 (2017).

87. Melzer TC, Cranston HJ, Weiss LM, Halonen SK. Host Cell Preference of Toxoplasma gondii Cysts in Murine Brain: A Confocal Study. J Neuroparasitology 1, (2010).

88. Rodriguez R, et al. Hepcidin induction by pathogens and pathogen-derived molecules is strongly dependent on interleukin-6. Infection and immunity 82, 745–752 (2014).

89. Chen AC, Donovan A, Ned-Sykes R, Andrews NC. Noncanonical role of transferrin receptor 1 is essential for intestinal homeostasis. Proceedings of the National Academy of Sciences of the United States of America 112, 11714–11719 (2015).

