## Supplemental figures and tables for "Theft of Host Transferrin Receptor-1 by *Toxoplasma gondii* is required for infection"

**Supplemental Table S1. Iron-Related Protein Homology Analysis in *Toxoplasma gondii***

Supplementary Table 1. Iron-Related Protein Homology Analysis in *Toxoplasma gondii*

| Iron Transport & Storage | | | | | | | | |
|---|---|---|---|---|---|---|---|---|
| **Protein** | **Reference Organism** | **Accession** | **Target ID** | **Bit score** | **E-value** | **Fitness Score (Sidik et al. 2016)** | **IFNg Fitness (Wang et al. 2020)** | **Reference** |
| DMT1 (Iso 2) | *Mus musculus* | NP_032758.2 | TGME49_267270 | 322 | 2.00E-95 | 0.49 | -0.38 | [1, 2] |
| NRAMP1 | *Mus musculus* | NP_038640.2 | TGME49_267270 | 306 | 2.00E-90 | 0.49 | -0.38 | [3] |
| VIT | *Plasmodium berghei* | PBANKA_143860 | TGME49_266800 | 255 | 6.00E-82 | -1.22 | 0.54 | [4] |
| YbtP | *Enterobacteria* | WP_001327262.1 | TGME49_269000 | 209 | 2.00E-56 | -4.96 | -0.09 | [5] |
| MntH | *Escherichia coli* | NP_416893.1 | TGME49_267270 | 179 | 2.00E-47 | 0.49 | -0.38 | [6] |
| DMT1 (Iso 3) | *Mus musculus* | NP_001343881.1 | TGME49_267270 | 162 | 5.00E-43 | 0.49 | -0.38 | [1, 2] |
| YbtQ | *Enterobacteria* | WP_012131848.1 | TGME49_273360 | 218 | 3.00E-39 | -0.17 | -0.24 | [5] |
| EntA | *Escherichia coli* | NP_415128.1 | TGME49_217740 | 103 | 2.00E-24 | -1.89 | -0.23 | [7, 8] |
| PchD | *Pseudomonas aeruginosa* | CAA57966.1 | TGME49_232580 | 240 | 4.00E-20 | -4.36 | -1.39 | [9] |
| FecE | *Escherichia coli* | AAA97183.1 | TGME49_269000 | 92.8 | 6.00E-20 | -4.96 | -0.09 | [10] |
| Irp9 | *Enterobacteria* | VTP73725.1 | TGME49_202920 | 235 | 6.00E-20 | -3.74 | -0.04 | [11] |
| YbtS | *Enterobacteria* | WP_012132137.1 | TGME49_202920 | 94.7 | 1.00E-19 | -3.74 | -0.04 | [12] |
| FhuC | *Escherichia coli* | CAD6022026.1 | TGME49_260310 | 90.1 | 1.00E-18 | 0.61 | 0.46 | [13] |
| YfeB | *Yersinia* | WP_002211843.1 | TGME49_273360 | 208 | 3.00E-17 | -0.17 | -0.24 | [6] |
| YbtE | *Enterobacteria* | WP_001088835.1 | TGME49_266640 | 80.9 | 5.00E-15 | 0.59 | -2.08 | [12] |
| PchA | *Pseudomonas aeruginosa* | CAA57969.1 | TGME49_202920 | 70.9 | 6.00E-12 | -3.74 | -0.04 | [14] |
| ZIPCO | *Plasmodium berghei* | PBANKA_0506500 | TGME49_261720 | 59.7 | 8.00E-09 | -4.31 | -0.03 | [15] |
| EntC | *Escherichia coli* | NP_415125.1 | TGME49_202920 | 54.7 | 5.00E-07 | -3.74 | -0.04 | [16] |
| ZIPCO | *Plasmodium berghei* | PBANKA_0506500 | TGME49_225530 | 52 | 3.00E-06 | -2.94 | -0.01 | [15] |
| AlcA | *Bordatella bronchiseptica* | AAB40618.1 | No Results | | | | | [17] |
| AlcB | *Bordatella bronchiseptica* | AAB40619.1 | No Results | | | | | [17] |
| AlcC | *Bordatella bronchiseptica* | AAB40620.1 | No Results | | | | | [17] |
| AlcD | *Bordatella pertussis* | WP_014905923.1 | No Results | | | | | [18] |
| AlcE | *Bordatella* | WP_005013753.1 | No Results | | | | | [19] |
| AlcS (Bcr) | *Bordatella* | WP_010930940.1 | No Results | | | | | [20] |
| Arn1 | *Saccharomyces cerevisiae* | NP_011823.1 | No Results | | | | | [21, 22] |

| | | | | | | | | | |
|---|---|---|---|---|---|---|---|---|---|
| Arn2 (Taf1) | *Saccharomyces cerevisiae* | NP_011816.2 | No Results | | | | | | [22, 23] |
| Arn3 (Sit1) | *Saccharomyces cerevisiae* | NP_010849.3 | No Results | | | | | | [22, 24] |
| Arn4 (ENB1) | *Saccharomyces cerevisiae* | NP_014484.1 | No Results | | | | | | [25] |
| Ceruloplasmin (ferroxidase) | *Homo sapiens* | ACB21047.1 | No Results | | | | | | [26] |
| CsbX | *Bacillus subtilis* | NP_390654.2 | No Results | | | | | | [27] |
| DMT1 (Iso 1) | *Mus musculus* | NP_001139633.1 | No Results | | | | | | [1, 2] |
| EfeB | *Escherichia coli* | NP_415538.1 | No Results | | | | | | [28] |
| EfeO | *Escherichia coli* | NP_415537.1 | No Results | | | | | | [28] |
| EfeU | *Escherichia coli* | AKK17243.1 | No Results | | | | | | [28] |
| EntB | *Escherichia coli* | NP_415127.1 | No Results | | | | | | [29] |
| EntS | *Escherichia coli* | NP_415123.1 | No Results | | | | | | [30] |
| Etf-3 (ECH0767) | *Ehrlichia chaffensis* | WP_006011519.1 | No Results | | | | | | [31] |
| FatA | *Vibrio anguillarum* | CDQ47873.1 | No Results | | | | | | [32] |
| FatB | *Vibrio anguillarum* | CDQ47872.1 | No Results | | | | | | [33] |
| FatC | *Vibrio anguillarum* | CDQ47871.1 | No Results | | | | | | [33] |
| FatD | *Vibrio anguillarum* | P37738.1 | No Results | | | | | | [33] |
| FauA | *Bordatella bronchiseptica* | WP_062758580.1 | No Results | | | | | | [34] |
| FecA | *Escherichia coli* | NP_418711.1 | No Results | | | | | | [10, 35] |
| FecB | *Escherichia coli* | NP_418710.4 | No Results | | | | | | [10, 36] |
| FecC | *Escherichia coli* | WP_000125187.1 | No Results | | | | | | [10, 36] |
| FecD | *Escherichia coli* | NP_418708.1 | No Results | | | | | | [10] |
| FeoA | *Escherichia coli* | NP_417867.1 | No Results | | | | | | [6] |
| FeoB | *Escherichia coli* | NP_417868.1 | No Results | | | | | | [6] |
| FeoC | *Escherichia coli* | NP_417869.1 | No Results | | | | | | [6, 37] |
| Ferroportin | *Mus musculus* | AAF80987.1 | No Results | | | | | | [38, 39] |
| FET4 | *Saccharomyces cerevisiae* | NP_014052.1 | No Results | | | | | | [40] |
| FeuA | *Bacillus subtilis* | NP_388044.1 | No Results | | | | | | [41] |
| FeuB | *Bacillus subtilis* | NP_388043.1 | No Results | | | | | | [42] |
| FeuC | *Bacillus subtilis* | NP_388042.2 | No Results | | | | | | [43] |
| FhuA | *Escherichia coli* | QIM05368.1 | No Results | | | | | | [44] |
| FhuB | *Escherichia coli* | CAD6022020.1 | No Results | | | | | | [44] |
| FhuD | *Escherichia coli* | CAD6017367.1 | No Results | | | | | | [44] |

| | | | | | | | | |
|---|---|---|---|---|---|---|---|---|
| FIT1 | *Saccharomyces cerevisiae* | NP_010823.1 | No Results | | | | | [45] |
| FIT2 | *Saccharomyces cerevisiae* | NP_015027.1 | No Results | | | | | [45] |
| FIT3 | *Saccharomyces cerevisiae* | NP_015028.3 | No Results | | | | | [45] |
| FptA | *Pseudomonas aeruginosa* | NP_252911.1 | No Results | | | | | [46] |
| FpvA | *Pseudomonas aeruginosa* | NP_251088.1 | No Results | | | | | [47] |
| FRO2 | *Arabidopsis thaliana* | CAA70770.1 | No Results | | | | | [48] |
| Ftn (Heavy) | *Homo sapiens* | NP_002023.2 | No Results | | | | | [49] |
| Ftn (Light) | *Homo sapiens* | NP_000137.2 | No Results | | | | | [49] |
| Ftr1 | *Saccharomyces cerevisiae* | NP_011072.1 | No Results | | | | | [50] |
| FyuA | *Enterobacteria* | WP_000784549.1 | No Results | | | | | [51] |
| HemO | *Leptospira interrogans* | AAN51745.1 | No Results | | | | | [52] |
| Hepcidin | *Homo sapiens* | AAH20612.1 | No Results | | | | | [53] |
| HFE | *Homo sapiens* | AAB82083.1 | No Results | | | | | [54] |
| IRT1 | *Arabidopsis thaliana* | NP_567590.3 | No Results | | | | | [55] |
| IRT2 | *Arabidopsis thaliana* | OAO98274.1 | No Results | | | | | [55] |
| IucA | *Escherichia coli* | CAA53707.1 | No Results | | | | | [56] |
| IucB | *Escherichia coli* | QBQ68952.1 | No Results | | | | | [56] |
| IucC | *Escherichia coli* | AAS66995.1 | No Results | | | | | [56] |
| IucD | *Escherichia coli* | QBQ68950.1 | No Results | | | | | [56] |
| LbpA | *Neisseria meningitidis* | WP_002229702.1 | No Results | | | | | [57] |
| LbpB | *Neisseria meningitidis* | WP_196979656.1 | No Results | | | | | [57] |
| LbtA | *Legionella pneumophila* | WP_015444551.1 | No Results | | | | | [58] |
| LbtB | *Legionella pneumophila* | WP_010947055.1 | No Results | | | | | [58] |
| LbtC | *Legionella pneumophila* | WP_010947054.1 | No Results | | | | | [59] |
| LbtU | *Legionella pneumophila* | P_010947057.1 | No Results | | | | | [60] |
| MexA | *Pseudomonas aeruginosa* | WP_049324879.1 | No Results | | | | | [61] |
| MexB | *Pseudomonas aeruginosa* | NP_249117.1 | No Results | | | | | [61] |
| OprM | *Pseudomonas aeruginosa* | NP_249118.1 | No Results | | | | | [61] |
| PchB | *Pseudomonas aeruginosa* | CAA57968.1 | No Results | | | | | [9] |
| PchC | *Pseudomonas aeruginosa* | CAA57967.1 | No Results | | | | | [62] |
| PfeA | *Pseudomonas aeruginosa* | NP_251378.1 | No Results | | | | | [63] |
| PupB | *Pseudomonas putida* | CAA51995.1 | No Results | | | | | [64] |
| PvsC | *Pseudomonas aeruginosa* | PTC36453.1 | No Results | | | | | [65] |

| | | | | | | | | |
|---|---|---|---|---|---|---|---|---|
| TbpA | *Neisseria meningitidis* | WP_004465694.1 | No Results | | | | | [66] |
| TbpB | *Neisseria gonorrhoeae* | STZ83427.1 | No Results | | | | | [66] |
| TfR2 | *Homo sapiens* | NP_003218.2 | No Results | | | | | [67] |
| TfRC | *Homo sapiens* | AAA61153.1 | No Results | | | | | [68] |
| TolC | *Escherichia coli* | NP_417507.2 | No Results | | | | | [69] |
| Transferrin | *Mus musculus* | NP_598738.1 | No Results | | | | | [68] |
| YbtT | *Enterobacteria* | WP_004175339.1 | No Results | | | | | [70] |
| YbtU | *Enterobacteria* | WP_012132135.1 | No Results | | | | | [70] |
| YbtX | *Enterobacteria* | WP_001286279.1 | No Results | | | | | [5, 71] |
| YfeA | *Yersinia* | WP_002211842.1 | No Results | | | | | [6] |
| YfeC | *Yersinia* | WP_004706545.1 | No Results | | | | | [6] |
| YfeD | *Yersinia* | WP_002211845.1 | No Results | | | | | [6] |
| YS1 | *Zea mays* | Q9AY27.1 | No Results | | | | | [6] |
| YSL1 | *Arabidopsis thaliana* | NP_567694.2 | No Results | | | | | [72] |
| YSL4 | *Arabidopsis thaliana* | NP_198916.2 | No Results | | | | | [73] |
| YSL6 | *Arabidopsis thaliana* | NP_566806.1 | No Results | | | | | [73] |
| ZupT | *Escherichia coli* | NP_417512.1 | No Results | | | | | [6] |

### Iron-Sulfur Complex Loading

| Protein | Reference Organism | Accession | Target ID | Bit score | E-value | Fitness Score (Sidik et al. 2016) | IFNg Fitness (Wang et al. 2020) | Reference |
|---|---|---|---|---|---|---|---|---|
| HscA | *Escherichia coli* | NP_417021.1 | TGME49_251780 | 425 | 1.00E-137 | -5.09 | -0.04 | [74] |
| NifS(NFS1) | *Salmonella enterica* | NP_461478.1 | TGME49_211090 | 413 | 3.00E-137 | -3.54 | 0.02 | [74, 75] |
| SufB | *Plasmodium berghei* | BAL70678.1 | TGME49_300650 | 395 | 2.00E-131 | | | [76, 77] |
| HscA | *Escherichia coli* | NP_417021.1 | TGME49_311720 | 399 | 2.00E-128 | -5.5 | -0.04 | [74] |
| SufC | *Plasmodium berghei* | SCM17963.1 | TGME49_225800 | 320 | 3.00E-103 | -5.57 | -0.04 | [76, 77] |
| SufD | *Plasmodium berghei* | SCO60652.1 | TGME49_273445 | 232 | 1.00E-60 | -5.29 | -0.89 | [76, 77] |
| SufS | *Plasmodium berghei* | SCO59130.1 | TGME49_216170 | 206 | 3.00E-56 | -3.95 | -0.74 | [76-78] |
| NifU | *Salmonella enterica* | NP_461477.1 | TGME49_237560 | 158 | 1.00E-47 | -5.99 | 0.17 | [74] |
| Fdx | *Escherichia coli* | NP_417020.1 | TGME49_240670 | 83.2 | 3.00E-19 | -2.45 | 0.04 | [74] |
| SufA | *Plasmodium berghei* | SCO62101.1 | TGME49_297925 | 87 | 1.00E-18 | -4.01 | 0.03 | [76, 77] |

| | | | | | | | | |
|---|---|---|---|---|---|---|---|---|
| IscA | *Escherichia coli* | NP_417023.1 | TGME49_297925 | 82 | 8.00E-18 | -4.01 | 0.03 | [74] |
| HscB | *Homo sapiens* | NP_741999.3 | TGME49_288685 | 84.7 | 1.00E-17 | -2.92 | -0.05 | [74] |
| SufE | *Plasmodium berghei* | SCM17131.1 | TGME49_277010 | 75.1 | 1.00E-14 | -3.31 | -0.11 | [76, 77] |
| CyaY (frataxin) | *Escherichia coli* | NP_418251.1 | TGME49_262810 | 54.7 | 9.00E-09 | -3.34 | -0.32 | [74] |
| SufU | *Bacillus subtilis* | QHE14930.1 | TGME49_237560 | 52 | 1.00E-07 | -5.99 | 0.17 | [76, 77] |
| NfuA | *Escherichia coli* | NP_417873.1 | TGME49_221922 | 46.6 | 4.00E-05 | -0.76 | -0.58 | [74] |
| CIA1 | *Toxoplasma gondii* | TGME49_313280 | N/A | N/A | N/A | -3.95 | -0.04 | [75] |
| CIA2 | *Toxoplasma gondii* | TGME49_306590 | N/A | N/A | N/A | -4.52 | -0.03 | [75] |
| Dre2 | *Toxoplasma gondii* | TGME49_216900 | N/A | N/A | N/A | -2.88 | -0.01 | [75] |
| IscU | *Toxoplasma gondii* | TGGT1_327560 | N/A | N/A | N/A | | | [78] |
| MMS19 | *Toxoplasma gondii* | TGME49_222230 | N/A | N/A | N/A | -5.93 | 0.02 | [75] |
| Nar1 | *Toxoplasma gondii* | TGME49_232580 | N/A | N/A | N/A | -4.36 | -1.39 | [75] |
| NBP35 | *Toxoplasma gondii* | TGME49_280730 | N/A | N/A | N/A | -4.81 | -0.01 | [75] |
| Tah18 | *Toxoplasma gondii* | TGME49_249320 | N/A | N/A | N/A | -3.87 | 0.01 | [75] |

#### Heme Biosynthesis

| Protein | Reference Organism | Accession | Target ID | Bit score | E-value | Fitness Score (Sidik et al. 2016) | IFNg Fitness (Wang et al. 2020) | Reference |
|---|---|---|---|---|---|---|---|---|
| ALAS | *Toxoplasma gondii* | TGME49_258690 | N/A | N/A | N/A | -4.58 | 0.49 | [79] |
| PBGS | *Toxoplasma gondii* | TGGT1_253900 | N/A | N/A | N/A | -4.41 | -0.34 | [79] |
| PBGD | *Toxoplasma gondii* | TGGT1_271420 | N/A | N/A | N/A | -3.93 | 0.06 | [79] |
| UROS | *Toxoplasma gondii* | TGGT1_264200 | N/A | N/A | N/A | -4.88 | -0.12 | [79] |
| UROD | *Toxoplasma gondii* | TGGT1_289940 | N/A | N/A | N/A | -4.6 | 0.01 | [79] |
| CPOX | *Toxoplasma gondii* | TGGT1_223020 | N/A | N/A | N/A | -4.64 | -0.02 | [79] |
| PPO | *Toxoplasma gondii* | TGGT1_272490 | N/A | N/A | N/A | -3.87 | -0.22 | [79] |
| FECH | *Toxoplasma gondii* | TGGT1_258650 | N/A | N/A | N/A | -5.69 | -0.29 | [79] |
| Tmem14c | *Toxoplasma gondii* | TGGT1_228110 | N/A | N/A | N/A | 1.29 | -0.34 | [80] |

#### Transcription / Translation

| Protein | Reference Organism | Accession | Target ID | Bit score | E-value | Fitness Score (Sidik et al. 2016) | IFNg Fitness (Wang et al. 2020) | Reference |
|---|---|---|---|---|---|---|---|---|
| Aconitase | *Mycobacterium tuberculosis* | O53166.1 | TGME49_226730 | 9421 | 0.00E+00 | -4.73 | -0.03 | [81] |
| IRP1 | *Homo sapiens* | NP_002188.1 | TGME49_226730 | 1069 | 0.00E+00 | -4.73 | -0.03 | [82] |
| IRP2 | *Homo sapiens* | AAI17484.1 | TGME49_226730 | 837 | 0.00E+00 | -4.73 | -0.03 | [83] |
| Myb2 | *Arabidopsis thaliana* | NP_182241.1 | TGME49_200385 | 85.5 | 4.00E-17 | 0.98 | -0.53 | [84, 85] |
| Myb3 | *Arabidopsis thaliana* | NP_564176.2 | TGME49_200385 | 82.8 | 2.00E-16 | 0.98 | -0.53 | [84-86] |
| AngR | *Vibrio anguillarum* | AAO92378.1 | TGME49_232580 | 186 | 5.00E-13 | 1.24 | -1.39 | [10] |
| Myb2 | *Arabidopsis thaliana* | NP_182241.1 | TGME49_275480 | 52 | 2.00E-06 | -4.24 | -0.04 | [84] |
| Myb3 | *Arabidopsis thaliana* | NP_564176.2 | TGME49_275480 | 52 | 2.00E-06 | -4.24 | -0.04 | [84, 86] |
| Aft1 | *Saccaromyces cerevisiae* | NP_011444.1 | No Results | | | | | [87] |
| Aft2 | *Saccaromyces cerevisiae* | KAF4002108.1 | No Results | | | | | [88] |
| AlcR | *Bordatella bronchiseptica* | WP_003814017.1 | No Results | | | | | [19] |
| AlgR | *Pseudomonas aeruginosa* | NP_253948.1 | No Results | | | | | [89] |
| AlgZ | *Pseudomonas aeruginosa* | NP_253949.1 | No Results | | | | | [89] |
| Btr | *Bacillus subtilis* | NP_388045.1 | No Results | | | | | [10] |
| DtxR | *Mycobacterium tuberculosis* | NP_217227.1 | No Results | | | | | [10] |
| FBXL5 | *Homo sapiens* | NP_036293.1 | No Results | | | | | [82] |
| FecI | *Escherichia coli* | NP_418713.1 | No Results | | | | | [10] |
| FecR | *Escherichia coli* | WP_001068910.1 | No Results | | | | | [10] |
| Fer | *Solanum lycopersicum* | NP_001234654.2 | No Results | | | | | [90] |
| FIT | *Arabidopsis thaliana* | Q0V7X4.1 | No Results | | | | | [91] |
| FIT | *Arabidopsis thaliana* | Q0V7X4.1 | No Results | | | | | [91] |
| FIT | *Arabidopsis thaliana* | Q0V7X4.1 | No Results | | | | | [91] |
| FpvI(PvdS) | *Pseudomonas aeruginosa* | NP_251077.1 | No Results | | | | | [10] |
| FpvR | *Pseudomonas aeruginosa* | NP_251078.1 | No Results | | | | | [92] |
| Fur | *Helicobacter pylori* | ABO20835.1 | No Results | | | | | [93] |
| Fur | *Escherichia coli* | AHG07399.1 | No Results | | | | | [10] |
| IrgB | *Vibrio cholerae* | AOY46299.1 | No Results | | | | | [10] |
| Myb1 | *Arabidopsis thaliana* | NP_187534.1 | No Results | | | | | [84] |
| Myb2 | *Arabidopsis thaliana* | NP_182241.1 | No Results | | | | | [84] |

| | | | | | | | | |
|---|---|---|---|---|---|---|---|---|
| PchA | *Pseudomonas aeruginosa* | AAA25926.1 | No Results | | | | | [10] |
| PfeR | *Pseudomonas aeruginosa* | NP_251376.1 | No Results | | | | | [10] |
| PfeS | *Pseudomonas aeruginosa* | NP_251377.1 | No Results | | | | | [10] |
| PupI | *Pseudomonas putida* | CAA54870.1 | No Results | | | | | [10] |
| PupR | *Pseudomonas putida* | CAA54871.1 | No Results | | | | | [94] |
| Sre | *Neurospora crassa* | Q1K8E7.1 | No Results | | | | | [95] |
| SreA | *Aspergillus nidulans* | AAD25328.1 | No Results | | | | | [96] |
| TonB | *Escherichia coli* | NP_415768.1 | No Results | | | | | [10] |
| Urbs1 | *Ustilago maydis* | AAB05617.1 | No Results | | | | | [97] |
| Yap5 | *Saccaromyces cerevisiae* | KZV10638.1 | No Results | | | | | [88] |
| YbtA | *Yersinia pestis* | WP_000140406.1 | No Results | | | | | [10] |

### Redox / Functionality

| Protein | Reference Organism | Accession | Target ID | Bit score | E-value | Fitness Score (Sidik et al. 2016) | IFNg Fitness (Wang et al. 2020) | Reference |
|---|---|---|---|---|---|---|---|---|
| COX1 | *Homo sapiens* | YP_003024028.1 | TGME49_255060 | 164 | 7.00E-44 | -1.95 | -0.11 | [98] |
| Cytochrome C | *Homo sapiens* | NP_061820.1 | TGME49_219750 | 137 | 5.00E-41 | -3.51 | -0.72 | [99] |
| eNOS (NOS3) Isoform 1 | *Homo sapiens* | NP_000594.2 | TGME49_219630 | 97.1 | 2.00E-19 | -2.78 | -0.5 | [100] |
| nNOS (NOS1) Isoform 2 | *Homo sapiens* | NP_001191142.1 | TGME49_219630 | 96.7 | 4.00E-19 | -2.78 | -0.5 | [101] |
| nNOS (NOS1) Isoform 1 | *Homo sapiens* | NP_000611.1 | TGME49_219630 | 96.3 | 3.00E-19 | -2.78 | -0.5 | [101] |
| nNOS (NOS1) Isoform 4 | *Homo sapiens* | NP_001191147.1 | TGME49_219630 | 96.7 | 4.00E-19 | -2.78 | -0.5 | [101] |
| ATPα | *Toxoplasma gondii* | TGME49_204400 | N/A | N/A | N/A | -3.84 | -0.41 | [102-104] |
| ATPβ | *Toxoplasma gondii* | TGME49_261950 | N/A | N/A | N/A | -4.84 | -0.04 | [102-104] |
| ATPγ | *Toxoplasma gondii* | TGME49_231910 | N/A | N/A | N/A | -3.94 | -0.04 | [102-104] |
| ATPδ | *Toxoplasma gondii* | TGME49_226000 | N/A | N/A | N/A | -4.57 | 0.76 | [102-104] |
| ATPε | *Toxoplasma gondii* | TGME49_314820 | N/A | N/A | N/A | -3.21 | -0.21 | [102-104] |
| ATP OSCP | *Toxoplasma gondii* | TGME49_284540 | N/A | N/A | N/A | -3.94 | -0.01 | [102-104] |
| ATPc | *Toxoplasma gondii* | TGME49_249720 | N/A | N/A | N/A | -2.98 | 0.1 | [102, 103] |

| | | | | | | | | |
|---|---|---|---|---|---|---|---|---|
| ATPb/ICAP2/ASAP-2 | *Toxoplasma gondii* | TGME49_231410 | N/A | N/A | N/A | -5.37 | -0.04 | [102-104] |
| ATPd/ICAP18/ASAP-3 | *Toxoplasma gondii* | TGME49_268830 | N/A | N/A | N/A | -2.02 | 0.13 | [102-104] |
| ATPk/ICAP6/ASAP-6 | *Toxoplasma gondii* | TGME49_260180 | N/A | N/A | N/A | -4.07 | 0.1 | [102-104] |
| ATPf/ICAP11/ASAP-1( | *Toxoplasma gondii* | TGME49_215610 | N/A | N/A | N/A | -3.55 | 0.16 | [102-104] |
| ATPTG3/ICAP8/ASAP | *Toxoplasma gondii* | TGME49_218940 | N/A | N/A | N/A | -3.92 | -0.04 | [102-104] |
| ATPTG6/ASAP-4 | *Toxoplasma gondii* | TGME49_223040 | N/A | N/A | N/A | -4.49 | -0.03 | [102-104] |
| ATPTG8/CHCH Domai | *Toxoplasma gondii* | TGME49_258060 | N/A | N/A | N/A | -4.07 | -0.04 | [102, 103] |
| ATPTG2/ICAP15/ASA | *Toxoplasma gondii* | TGME49_282180 | N/A | N/A | N/A | -2.46 | 0.18 | [102-104] |
| ATPTG9/ASAP-9/CHC | *Toxoplasma gondii* | TGME49_285510 | N/A | N/A | N/A | -1.87 | -0.09 | [102-104] |
| ATPa/ASAP-1 | *Toxoplasma gondii* | TGME49_310360 | N/A | N/A | N/A | -4.49 | -0.04 | [102-104] |
| ATPTG15/ICAP9/ASA | *Toxoplasma gondii* | TGME49_247410 | N/A | N/A | N/A | -3.9 | -0.63 | [102, 104] |
| IF1 | *Toxoplasma gondii* | TGME49_215350 | N/A | N/A | N/A | 1.82 | -1.01 | [102-104] |
| RPS11 | *Toxoplasma gondii* | TGME49_226970 | N/A | N/A | N/A | -3.75 | -0.05 | [102] |
| ApiCox25 | *Toxoplasma gondii* | TGGT1_264040 | N/A | N/A | N/A | -2.54 | 0.52 | [105] |
| ApiCox16 | *Toxoplasma gondii* | TGGT1_265370 | N/A | N/A | N/A | 1.56 | -0.44 | [105] |
| ApiCox5b | *Toxoplasma gondii* | TGGT1_209260 | N/A | N/A | N/A | -3.07 | -0.09 | [105] |
| ApiCox18 | *Toxoplasma gondii* | TGGT1_221510 | N/A | N/A | N/A | -3.28 | 0.01 | [105] |
| ApiCox23 | *Toxoplasma gondii* | TGGT1_262640 | N/A | N/A | N/A | -3.49 | -0.33 | [105] |
| ApiCox30 | *Toxoplasma gondii* | TGGT1_297810 | N/A | N/A | N/A | -3.64 | -0.17 | [105] |
| ApiCox19 | *Toxoplasma gondii* | TGGT1_247770 | N/A | N/A | N/A | -2.61 | -1.27 | [105] |
| ApiCox35 | *Toxoplasma gondii* | TGGT1_229920 | N/A | N/A | N/A | -3.84 | -0.03 | [105] |
| ApiCox26 | *Toxoplasma gondii* | TGGT1_306670 | N/A | N/A | N/A | -3.68 | -0.18 | [105] |
| ApiCox2a | *Toxoplasma gondii* | TGGT1_226590 | N/A | N/A | N/A | -3.8 | -0.69 | [105] |
| ApiCox2b | *Toxoplasma gondii* | TGGT1_310470 | N/A | N/A | N/A | -4.18 | -0.38 | [105] |
| ApiCox24 | *Toxoplasma gondii* | TGGT1_286530 | N/A | N/A | N/A | -2.82 | -0.3 | [105] |
| ApiCox13 | *Toxoplasma gondii* | TGGT1_254030 | N/A | N/A | N/A | -4.26 | -0.04 | [105] |
| ApiCox14 | *Toxoplasma gondii* | TGGT1_242840 | N/A | N/A | N/A | -3.58 | -0.04 | [105] |
| Cox3 | *Toxoplasma gondii* | TGVEG_442760 | N/A | N/A | N/A | | | [105] |
| ATPij/ASAP-11 | *Toxoplasma gondii* | TGME49_290030 | N/A | N/A | N/A | -3.88 | -0.05 | [103, 104] |
| ATPTG17/ASAP-12 | *Toxoplasma gondii* | TGME49_310180 | N/A | N/A | N/A | -3.4 | -0.04 | [103, 104] |
| ATPG10/ASAP-13 | *Toxoplasma gondii* | TGME49_214930 | N/A | N/A | N/A | -1.37 | -0.15 | [103, 104] |
| ATPTG12/ASAP-14 | *Toxoplasma gondii* | TGME49_245450 | N/A | N/A | N/A | -2.95 | -0.08 | [103, 104] |
| ATP8/ASAP-15 | *Toxoplasma gondii* | TGME49_208440 | N/A | N/A | N/A | -3.54 | -0.33 | [103, 104] |

| | | | | | | | | |
|---|---|---|---|---|---|---|---|---|
| ATPTG4/ASAP-16 | *Toxoplasma gondii* | TGME49_201800 | N/A | N/A | N/A | -4.01 | 0.12 | [103, 104] |
| ATPTG13/ASAP-17 | *Toxoplasma gondii* | TGME49_225730 | N/A | N/A | N/A | -3.65 | -0.25 | [103, 104] |
| ATPTG14/ASAP-18 | *Toxoplasma gondii* | TGME49_263080 | N/A | N/A | N/A | -3.1 | 0.06 | [103, 104] |
| ATPTG11/ASAP-19 | *Toxoplasma gondii* | TGME49_263990 | N/A | N/A | N/A | 0.22 | -0.13 | [103, 104] |
| ATPTG5/ASAP-20 | *Toxoplasma gondii* | TGME49_270360 | N/A | N/A | N/A | 0.32 | -0.24 | [103, 104] |
| ATPTG1 | *Toxoplasma gondii* | TGGT1_246540 | N/A | N/A | N/A | -4.36 | 0.04 | [103, 104] |
| ATPTG7 | *Toxoplasma gondii* | TGGT1_290710 | N/A | N/A | N/A | 0.67 | -0.41 | [103, 104] |
| ATPTG16 | *Toxoplasma gondii* | TGGT1_211060 | N/A | N/A | N/A | -3.31 | 0.26 | [103, 104] |
| Catalase | *Toxoplasma gondii* | TGME49_232250 | N/A | N/A | N/A | 2.08 | -0.25 | [106] |
| Cytochrome p450 | *Toxoplasma gondii* | TGGT1_315770 | N/A | N/A | N/A | 0.80 | -1.08 | [107] |
| SOD | *Toxoplasma gondii* | TGME49_316310 | N/A | N/A | N/A | -4.42 | -0.03 | [108] |
| SOD2 | *Toxoplasma gondii* | TGME49_316330 | N/A | N/A | N/A | -4.09 | -0.04 | [109] |
| SOD3 | *Toxoplasma gondii* | TGME49_316190 | N/A | N/A | N/A | 0.26 | -0.09 | |
| Ceruloplasmin | *Homo sapiens* | XP_006713562.1 | No Results | | | | | [26] |
| COX4 | *Homo sapiens* | NP_001852.1 | No Results | | | | | [99] |
| Duodenal cytochrome B reductase | *Mus musculus* | NP_082869.2 | No Results | | | | | [110] |
| eNOS (NOS3) Isoform 2 | *Homo sapiens* | NP_001153581.1 | No Results | | | | | [100] |
| eNOS (NOS3) Isoform 3 | *Homo sapiens* | NP_001153582.1 | No Results | | | | | [100] |
| eNOS (NOS3) Isoform 4 | *Homo sapiens* | NP_001153583.1 | No Results | | | | | [100] |
| FET3 | *Saccharomyces cerevisiae* | NP_013774.1 | No Results | | | | | [111] |
| Fre1 | *Saccharomyces cerevisiae* | NP_013315.1 | No Results | | | | | [22, 112] |
| Fre2 | *Saccharomyces cerevisiae* | CAA82065.1 | No Results | | | | | [22] |
| Fre3 | *Saccharomyces cerevisiae* | NP_015026.1 | No Results | | | | | [22] |
| Fre4 | *Saccharomyces cerevisiae* | NP_014458.1 | No Results | | | | | [22] |
| Fre5 | *Saccharomyces cerevisiae* | NP_015029.1 | No Results | | | | | [22] |
| Fre6 | *Saccharomyces cerevisiae* | NP_013049.1 | No Results | | | | | [22] |
| Hephaestin | *Homo sapiens* | AAK08131.1 | No Results | | | | | [39, 113, 114] |
| HMOX-1 | *Homo sapiens* | NP_002124.1 | No Results | | | | | [115] |
| iNOS Chain A | *Mus musculus* | 2ORQ_A | No Results | | | | | [100] |

| | | | | | | | | |
|---|---|---|---|---|---|---|---|---|
| iNOS Chain B | *Mus musculus* | 1JWK_B | No Results | | | | | [100] |
| Steap1 | *Homo sapiens* | NP_036581.1 | No Results | | | | | [116] |
| Steap2 | *Homo sapiens* | NP_001231873.1 | No Results | | | | | [116] |
| Steap3 | *Homo sapiens* | NP_878919.2 | No Results | | | | | [112] |
| Steap4 | *Homo sapiens* | NP_078912.2 | No Results | | | | | [116] |

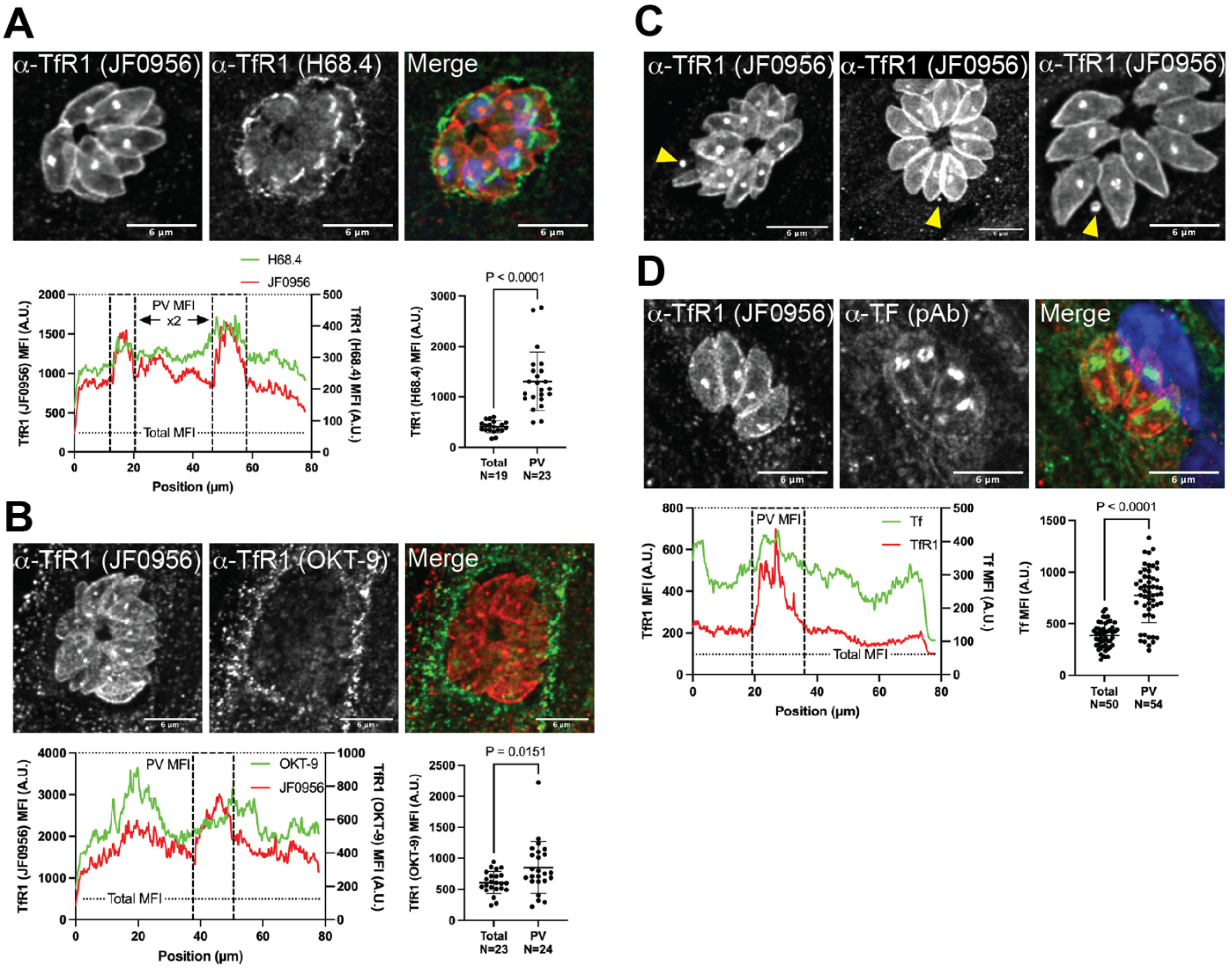


**Extended Data 1. Host transferrin receptor 1 and cognate ligand transferrin are acquired by *T. gondii*.**

RH-strain *T. gondii* 24-hour infections of MRC5 cells were stained for confocal IFA of host TfR1 and Tf. **A)** Representative IFA and quantitation of parasite associated TfR1 comparing JF0956 and H68.4 clones. Data pooled from two technical repeats and analyzed by unpaired student's t-test (t=6.648, df=40) **B)** Representative IFA and quantitation of parasite-associated TfR1 comparing JF0956 and OKT-9 clones. Data pooled from two technical repeats and analyzed by unpaired student's t-test (t=2.526, df=45) **C)** Chance observations of parasite-adjacent TfR1 (JF0956) puncta. **D)**

Representative IFA and quantitation of parasite associated TfR1 (JF0956) and Tf (polyclonal). Data pooled from two technical repeats and analyzed by unpaired student's t-test (t=9.462, df=102).

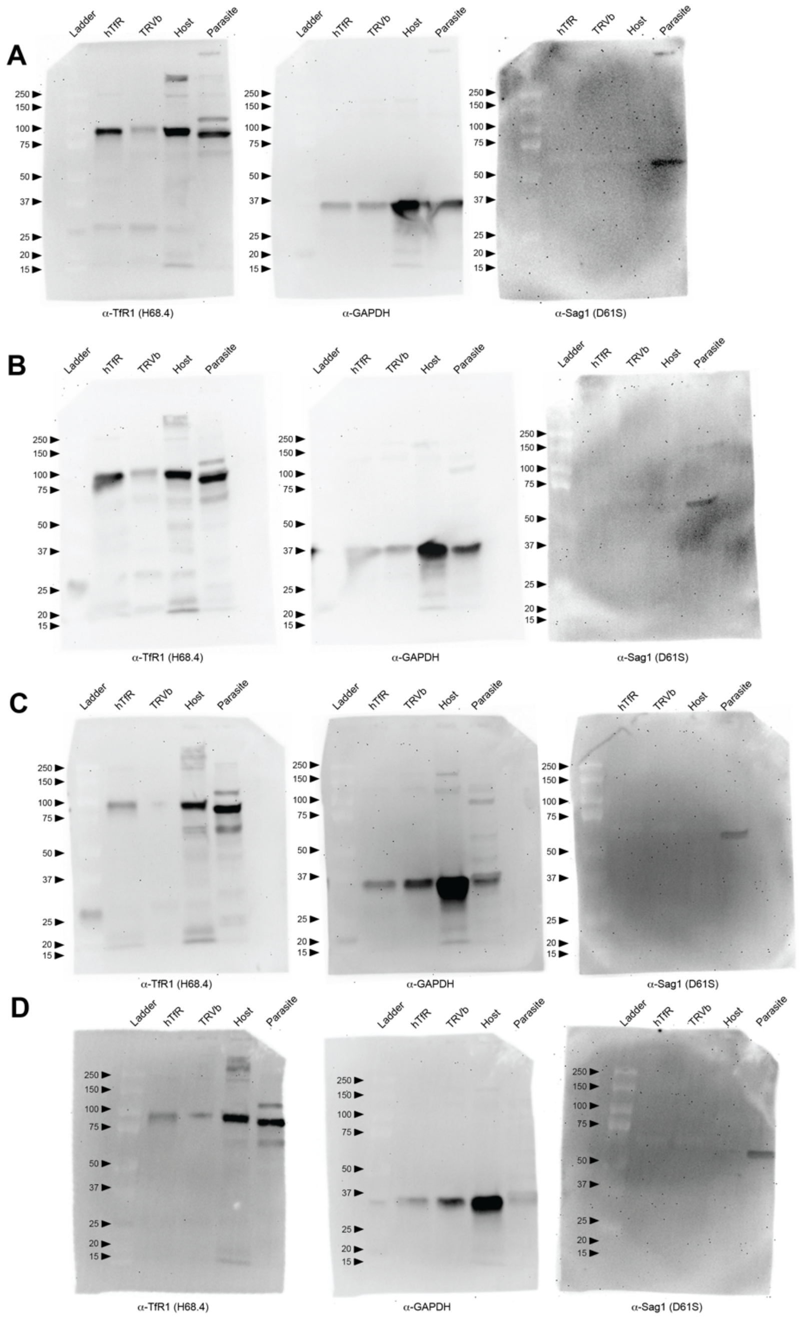


**Extended Data 2. Full TfR1 Western Blot replicates.**

30μg of cell lysates were probed in Western Blot for TfR1. Positive control hTfR cell lysates are TfR1-deficient cells reconstituted with the human TfR1 gene. Negative control TRVb cells are the TfR1-deficient cell. The uninfected control is host cell lysate from uninfected MRC5 cells. Purified parasites are RH strain *T. gondii* mechanically lysed from host cells and filter purified. **A)** Experiment 1 membrane stripped and reprobed for TfR1 (H68.4), GAPDH, and Sag1. **B)** Experiment 2 membrane stripped and reprobed for TfR1 (H68.4), GAPDH, and Sag1. **C)** Experiment 3 membrane stripped and reprobed for TfR1 (H68.4), GAPDH, and Sag1. **D)** Experiment 4 membrane stripped and reprobed for TfR1 (H68.4), GAPDH, and Sag1.

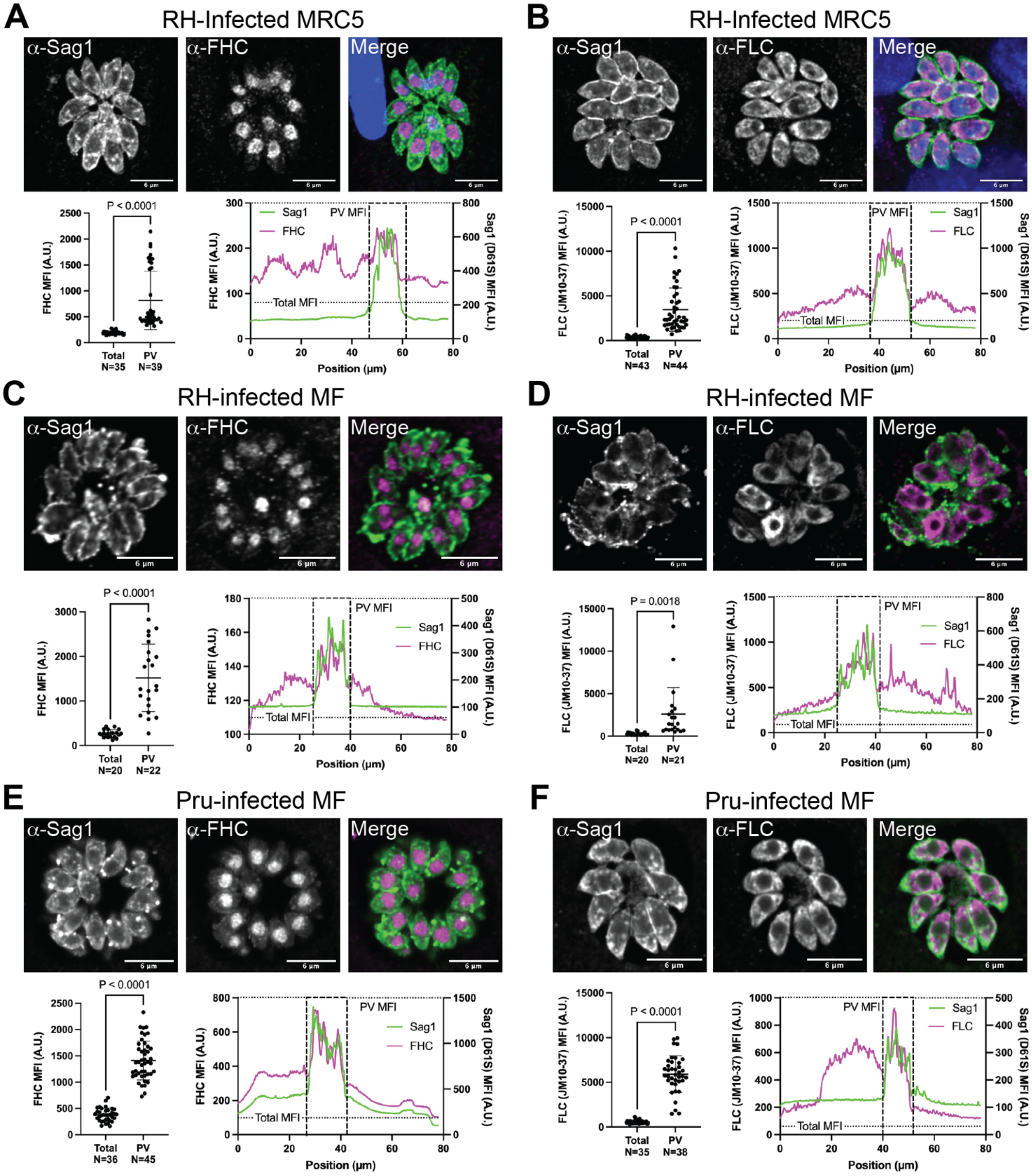


**Extended Data 3. Ferritin Heavy and Light Chains are incorporation occurs irrespective of virulence type or host cell genus.**

Shown are representative images, distribution plot, and quantitation of parasite associated: **A)** FHC of RH-strain *T. gondii* infecting human cells. Data pooled from four technical repeats and analyzed by unpaired student's t-test (t=6.485, df=72). **B)** FLC of RH-strain *T. gondii* infecting human cells. Data pooled from four technical repeats and analyzed by unpaired student's t-test (t=8.387, df=85). **C)** FHC of RH-strain *T. gondii* infecting MF cells. Data pooled from two technical repeats and analyzed by unpaired student's t-test (t=7.350, df=40). **D)** FLC of RH-strain *T. gondii* infecting MF cells. Data pooled from two technical repeats and analyzed by unpaired student's t-test (t=3.346, df=39). **E)** FHC of Pru-strain *T. gondii* infecting MF cells. Data pooled from two technical repeats and analyzed by unpaired student's t-test (t=15.95, df=79). **F)** FLC of Pru-strain *T. gondii* infecting MF cells. Data pooled from two technical repeats and analyzed with unpaired student's t-test (t=15.32, df=71).

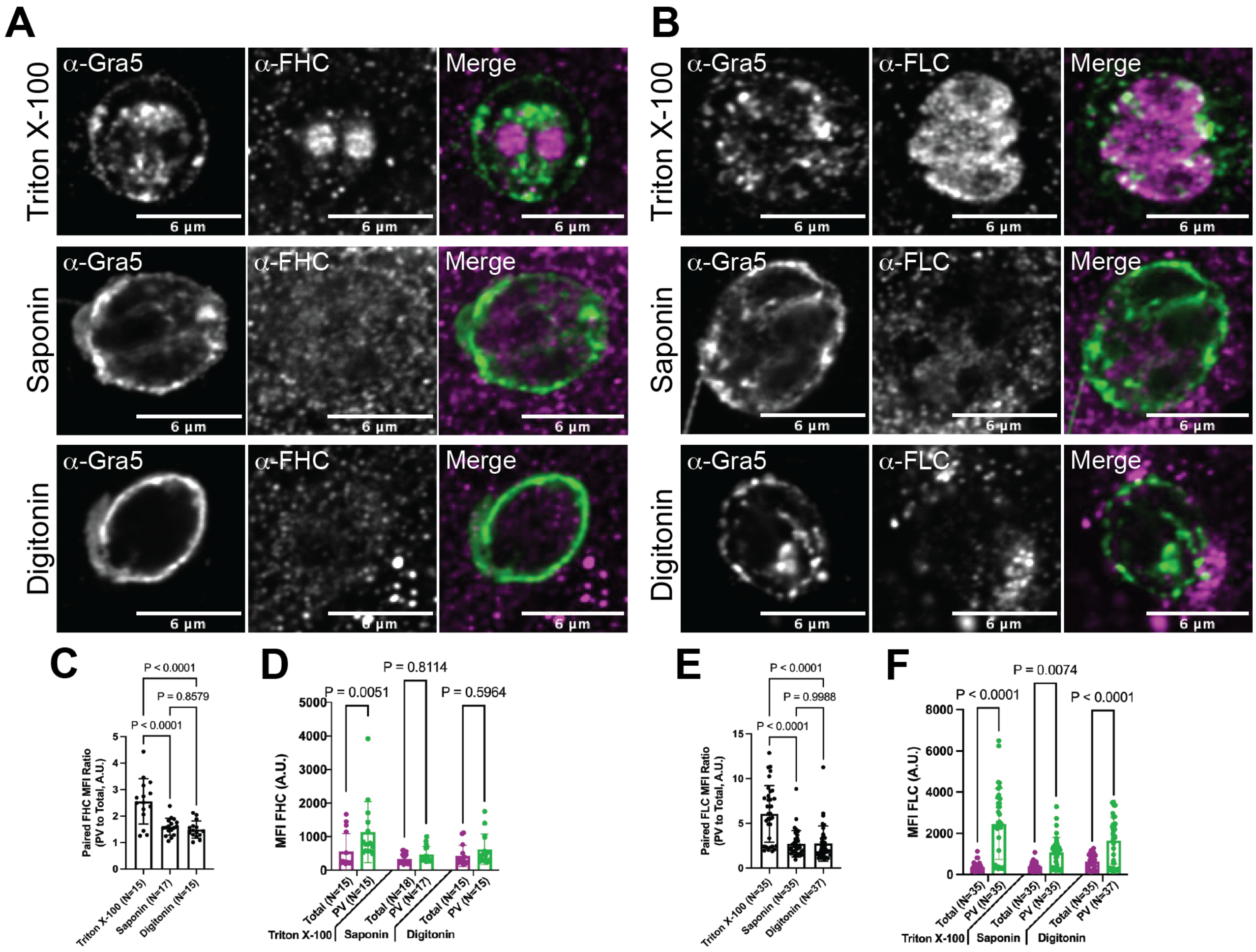


**Extended Data 4. Ferritin Heavy and Light Chains are incorporated into the parasite body.**

RH-strain *T. gondii* 24-hour infections of MRC5 cells were differentially permeabilized and stained for confocal IFA of host FHC and FLC. **A)** Representative FHC-IFA images of samples permeabilized with Triton X-100 (Top Row), Saponin (Middle Row) or Digitonin (Bottom Row). **B)** Representative FLC-IFA images of samples permeabilized with Triton X-100 (Top Row), Saponin (Middle Row) or Digitonin (Bottom Row). **C)** Comparison of PV/Total FHC MFI ratio between permeabilization type. Analysis performed by Ordinary One-Way ANOVA (f=17.17, df=46). **D)** Permeabilization-grouped comparisons of Total and PV-associated FHC. Analysis performed by Two-Way ANOVA (f=6.846, df=40). **E)** Comparison of PV/Total FLC MFI ratio between permeabilization type. Analysis performed by Ordinary One-Way ANOVA (f=24.55, df=106). **F)** Permeabilization-grouped comparisons of Total and PV-associated FLC. Analysis performed by Two-Way ANOVA (f=9.706, df=206). All data pooled from two technical repeats.

| Iron Transport & Storage - *Plasmodium* | | | | | |
|---|---|---|---|---|---|
| **Protein** | **Reference Organism** | **Accession** | ***P. falciparum* Target ID** | **Bit score** | **E-value** |
| DMT1 | *Mus musculus* | NP_032758.2 | PF3D7_0523800 | 185 | 3.00E-51 |
| NRAMP1 | *Mus musculus* | NP_038640.2 | PF3D7_0523800 | 168 | 2.00E-45 |
| Ferroportin | *Mus musculus* | AAI50987.1 | No Results | - | - |
| Ftn (Heavy) | *Homo sapiens* | NP_002023.2 | No Results | - | - |
| Ftn (Light) | *Homo sapiens* | NP_000137.2 | No Results | - | - |
| Hepcidin | *Homo sapiens* | AAH20612.1 | No Results | - | - |
| HFE | *Homo sapiens* | AAB82083.1 | No Results | - | - |
| TfR2 | *Homo sapiens* | NP_003218.2 | No Results | - | - |
| TfRC | *Homo sapiens* | AAA61153.1 | No Results | - | - |
| Transferrin | *Mus musculus* | NP_598738.1 | No Results | - | - |

**Supplemental Table 2.** ***Plasmodium falciparum*** **homology to mammalian iron transport and storage proteins.**

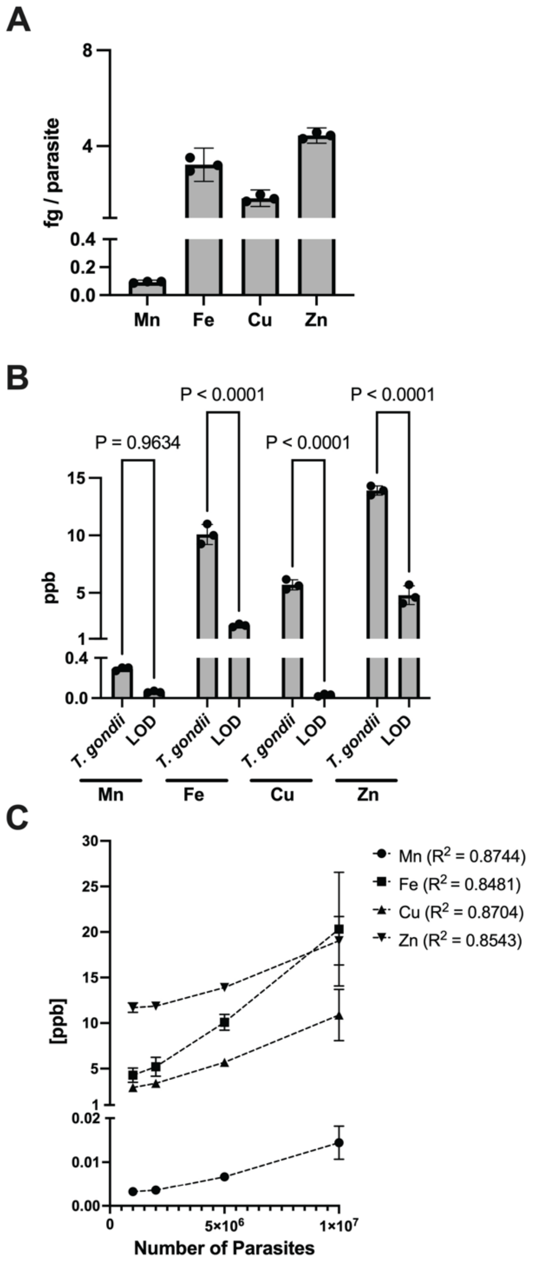


**Supplemental Figure 5: Proof-of-concept of ICPMS elemental analysis.**

RH strain *T. gondii* was isolated from routine passage in MRC5 cells. Purified parasites were digested for elemental composition analysis by ICPMS at sample concentrations indicated. Depicted are the mass of each element on a per cell basis. **A)** Measured mass per cell of manganese, iron, copper, and zinc at five million parasites per sample. **B)** Comparisons of each element to the limit of detection (LOD) for this method at five million parasites per sample. Analysis performed by Two-Way ANOVA (f=873.2, df=16). **C)** Measurement of total elemental content dependent on cell concentration per sample. Experiment performed once as a proof of concept.

**Movie 1. Theft of Transferrin Receptor 1 by *T. gondii***

RH-strain *T. gondii* infecting MRC5 fibroblasts for 24 hours was IFA stained for TfR1, Sag1, and DAPI. Z-stack images were taken in 0.05μm intervals. Movie depicts the 3D-projection where TfR1 fluorescence is shown in red, Sag1 fluorescence is shown in green, and DAPI is shown in blue.

**Movie 2. Theft of Transferrin by *T. gondii***

RH-strain *T. gondii* infecting MRC5 fibroblasts for 24 hours was IFA stained for TfR1, Tf, and DAPI. Z-stack images were taken in 0.05μm intervals. Movie depicts the 3D-projection where TfR1 fluorescence is shown in red, Tf fluorescence is shown in green, and DAPI is shown in blue.

**Movie 3. Inheritance of host TfR1 by *T. gondii* daughter cells during endodyogeny**

RH-strain *T. gondii* infecting MRC5 fibroblasts for 24 hours was IFA stained for TfR1, Sag1, and DAPI. Z-stack images were taken in 0.05μm intervals. Movie depicts the 3D-projection where TfR1 fluorescence is shown in red and Sag1 fluorescence is shown in green.
