## Supplementary material for "Theft of Host Transferrin Receptor-1 by *Toxoplasma gondii* is required for infection": References for suplemental table 1

1.	Fleming, M.D., et al., *Microcytic anaemia mice have a mutation in Nramp2, a candidate iron transporter gene.* Nat Genet, 1997. **16**(4): p. 383-6.

2.	Gunshin, H., et al., *Cloning and characterization of a mammalian proton-coupled metal-ion transporter.* Nature, 1997. **388**(6641): p. 482-8.

3.	Soe-Lin, S., et al., *Nramp1 promotes efficient macrophage recycling of iron following erythrophagocytosis in vivo.* Proc Natl Acad Sci U S A, 2009. **106**(14): p. 5960-5.

4.	Slavic, K., et al., *A vacuolar iron-transporter homologue acts as a detoxifier in Plasmodium.* 2016. **7**: p. 10403.

5.	Fetherston, J.D., V.J. Bertolino, and R.D. Perry, *YbtP and YbtQ: two ABC transporters required for iron uptake in Yersinia pestis.* Mol Microbiol, 1999. **32**(2): p. 289-99.

6.	Lau, C.K.Y., K.D. Krewulak, and H.J. Vogel, *Bacterial ferrous iron transport: the Feo system.* FEMS Microbiology Reviews, 2016. **40**(2): p. 273-298.

7.	Liu, J., K. Duncan, and C.T. Walsh, *Nucleotide sequence of a cluster of Escherichia coli enterobactin biosynthesis genes: identification of entA and purification of its product 2,3-dihydro-2,3-dihydroxybenzoate dehydrogenase.* J Bacteriol, 1989. **171**(2): p. 791-8.

8.	Nahlik, M.S., et al., *Nucleotide sequence and transcriptional organization of the Escherichia coli enterobactin biosynthesis cistrons entB and entA.* J Bacteriol, 1989. **171**(2): p. 784-90.

9.	Serino, L., et al., *Biosynthesis of pyochelin and dihydroaeruginoic acid requires the iron-regulated pchDCBA operon in Pseudomonas aeruginosa.* J Bacteriol, 1997. **179**(1): p. 248-57.

10.	Miethke, M. and M.A. Marahiel, *Siderophore-based iron acquisition and pathogen control.* Microbiol Mol Biol Rev, 2007. **71**(3): p. 413-51.

11.	Pelludat, C., D. Brem, and J. Heesemann, *Irp9, encoded by the high-pathogenicity island of Yersinia enterocolitica, is able to convert chorismate into salicylate, the precursor of the siderophore yersiniabactin.* J Bacteriol, 2003. **185**(18): p. 5648-53.

12.	Gehring, A.M., et al., *Iron acquisition in plague: modular logic in enzymatic biogenesis of yersiniabactin by Yersinia pestis.* Chem Biol, 1998. **5**(10): p. 573-86.

13.	Wooldridge, K.G., J.A. Morrissey, and P.H. Williams, *Transport of ferric-aerobactin into the periplasm and cytoplasm of Escherichia coli K12: role of envelope-associated proteins and effect of endogenous siderophores.* J Gen Microbiol, 1992. **138**(3): p. 597-603.

14.	Gaille, C., C. Reimmann, and D. Haas, *Isochorismate synthase (PchA), the first and rate-limiting enzyme in salicylate biosynthesis of Pseudomonas aeruginosa.* J Biol Chem, 2003. **278**(19): p. 16893-8.

15.	Sahu, T., et al., *ZIPCO, a putative metal ion transporter, is crucial for Plasmodium liver-stage development.* EMBO Molecular Medicine, 2014. **6**(11): p. 1387-1397.

16.	Liu, J., et al., *Overexpression, purification, and characterization of isochorismate synthase (EntC), the first enzyme involved in the biosynthesis of enterobactin from chorismate.* Biochemistry, 1990. **29**(6): p. 1417-25.

17.	Kang, H.Y., et al., *Identification and characterization of iron-regulated Bordetella pertussis alcaligin siderophore biosynthesis genes.* J Bacteriol, 1996. **178**(16): p. 4877-84.

18.	Funahashi, T., et al., *An iron-regulated gene required for utilization of aerobactin as an exogenous siderophore in Vibrio parahaemolyticus.* Microbiology (Reading), 2003. **149**(Pt 5): p. 1217-1225.

19.	Pradel, E., N. Guiso, and C. Locht, *Identification of AlcR, an AraC-type regulator of alcaligin siderophore synthesis in Bordetella bronchiseptica and Bordetella pertussis.* J Bacteriol, 1998. **180**(4): p. 871-80.

20.	Brickman, T.J. and S.K. Armstrong, *Bordetella AlcS transporter functions in alcaligin siderophore export and is central to inducer sensing in positive regulation of alcaligin system gene expression.* J Bacteriol, 2005. **187**(11): p. 3650-61.

21.	Heymann, P., J.F. Ernst, and G. Winkelmann, *Identification and substrate specificity of a ferrichrome-type siderophore transporter (Arn1p) in Saccharomyces cerevisiae.* FEMS Microbiol Lett, 2000. **186**(2): p. 221-7.

22.	Yun, C.W., et al., *Siderophore-iron uptake in saccharomyces cerevisiae. Identification of ferrichrome and fusarinine transporters.* J Biol Chem, 2000. **275**(21): p. 16354-9.

23.	Heymann, P., J.F. Ernst, and G. Winkelmann, *Identification of a fungal triacetylfusarinine C siderophore transport gene (TAF1) in Saccharomyces cerevisiae as a member of the major facilitator superfamily.* Biometals, 1999. **12**(4): p. 301-6.

24.	Lesuisse, E., M. Simon-Casteras, and P. Labbe, *Siderophore-mediated iron uptake in Saccharomyces cerevisiae: the SIT1 gene encodes a ferrioxamine B permease that belongs to the major facilitator superfamily.* Microbiology (Reading), 1998. **144 ( Pt 12)**: p. 3455-3462.

25.	Froissard, M., et al., *Trafficking of siderophore transporters in Saccharomyces cerevisiae and intracellular fate of ferrioxamine B conjugates.* Traffic, 2007. **8**(11): p. 1601-16.

26.	Van Eden, M.E. and S.D. Aust, *Intact human ceruloplasmin is required for the incorporation of iron into human ferritin.* Arch Biochem Biophys, 2000. **381**(1): p. 119-26.

27.	Page, W.J., et al., *The csbX gene of Azotobacter vinelandii encodes an MFS efflux pump required for catecholate siderophore export.* FEMS Microbiol Lett, 2003. **228**(2): p. 211-6.

28.	Cao, J., et al., *EfeUOB (YcdNOB) is a tripartite, acid-induced and CpxAR-regulated, low-pH Fe2+ transporter that is cryptic in Escherichia coli K-12 but functional in E. coli O157:H7.* Mol Microbiol, 2007. **65**(4): p. 857-75.

29.	Gehring, A.M., K.A. Bradley, and C.T. Walsh, *Enterobactin biosynthesis in Escherichia coli: isochorismate lyase (EntB) is a bifunctional enzyme that is phosphopantetheinylated by EntD and then acylated by EntE using ATP and 2,3-dihydroxybenzoate.* Biochemistry, 1997. **36**(28): p. 8495-503.

30.	Holden, V.I., et al., *Iron Acquisition and Siderophore Release by Carbapenem-Resistant Sequence Type 258 Klebsiella pneumoniae.* mSphere, 2018. **3**(2).

31.	Yan, Q., et al., *Iron robbery by intracellular pathogen via bacterial effector-induced ferritinophagy.* Proceedings of the National Academy of Sciences of the United States of America, 2021. **118**(23).

32.	Köster, W.L., et al., *Molecular characterization of the iron transport system mediated by the pJM1 plasmid in Vibrio anguillarum 775.* J Biol Chem, 1991. **266**(35): p. 23829-33.

33.	Naka, H., C.S. López, and J.H. Crosa, *Role of the pJM1 plasmid-encoded transport proteins FatB, C and D in ferric anguibactin uptake in the fish pathogen Vibrio anguillarum.* Environ Microbiol Rep, 2010. **2**(1): p. 104-111.

34.	Brickman, T.J. and S.K. Armstrong, *Essential role of the iron-regulated outer membrane receptor FauA in alcaligin siderophore-mediated iron uptake in Bordetella species.* J Bacteriol, 1999. **181**(19): p. 5958-66.

35.	Chakraborty, R., E. Storey, and D. van der Helm, *Molecular mechanism of ferricsiderophore passage through the outer membrane receptor proteins of Escherichia coli.* Biometals, 2007. **20**(3-4): p. 263-74.

36.	Braun, V. and F. Endriss, *Energy-coupled outer membrane transport proteins and regulatory proteins.* Biometals, 2007. **20**(3-4): p. 219-31.

37.	Smith, A.T., et al., *The FeoC [4Fe-4S] Cluster Is Redox-Active and Rapidly Oxygen-Sensitive.* Biochemistry, 2019. **58**(49): p. 4935-4949.

38.	Abboud, S. and D.J. Haile, *A novel mammalian iron-regulated protein involved in intracellular iron metabolism.* J Biol Chem, 2000. **275**(26): p. 19906-12.

39.	Han, O. and E.Y. Kim, *Colocalization of ferroportin-1 with hephaestin on the basolateral membrane of human intestinal absorptive cells.* J Cell Biochem, 2007. **101**(4): p. 1000-10.

40.	Dix, D., et al., *Characterization of the FET4 protein of yeast. Evidence for a direct role in the transport of iron.* J Biol Chem, 1997. **272**(18): p. 11770-7.

41.	Peuckert, F., et al., *The siderophore binding protein FeuA shows limited promiscuity toward exogenous triscatecholates.* Chem Biol, 2011. **18**(7): p. 907-19.

42.	Ichihara, S. and S. Mizushima, *Identification of an outer membrane protein responsible for the binding of the Fe-enterochelin complex to Escherichia coli cells.* J Biochem, 1978. **83**(1): p. 137-40.

43.	Urzúa, L.S., et al., *Identification and characterization of an iron ABC transporter operon in Gluconacetobacter diazotrophicus Pal 5.* Arch Microbiol, 2013. **195**(6): p. 431-8.

44.	Fecker, L. and V. Braun, *Cloning and expression of the fhu genes involved in iron(III)-hydroxamate uptake by Escherichia coli.* J Bacteriol, 1983. **156**(3): p. 1301-14.

45.	Protchenko, O., et al., *Three cell wall mannoproteins facilitate the uptake of iron in Saccharomyces cerevisiae.* J Biol Chem, 2001. **276**(52): p. 49244-50.

46.	Hoegy, F., et al., *Binding of iron-free siderophore, a common feature of siderophore outer membrane transporters of Escherichia coli and Pseudomonas aeruginosa.* J Biol Chem, 2005. **280**(21): p. 20222-30.

47.	Nader, M., et al., *Mechanism of ferripyoverdine uptake by Pseudomonas aeruginosa outer membrane transporter FpvA: no diffusion channel formed at any time during ferrisiderophore uptake.* Biochemistry, 2011. **50**(13): p. 2530-40.

48.	Kim, S.A., et al., *The iron deficiency response in Arabidopsis thaliana requires the phosphorylated transcription factor URI.* Proc Natl Acad Sci U S A, 2019. **116**(50): p. 24933-24942.

49.	Harrison, P.M. and P. Arosio, *The ferritins: molecular properties, iron storage function and cellular regulation.* Biochimica et Biophysica Acta (BBA) - Bioenergetics, 1996. **1275**(3): p. 161-203.

50.	Askwith, C. and J. Kaplan, *An oxidase-permease-based iron transport system in Schizosaccharomyces pombe and its expression in Saccharomyces cerevisiae.* J Biol Chem, 1997. **272**(1): p. 401-5.

51.	Haag, H., et al., *Purification of yersiniabactin: a siderophore and possible virulence factor of Yersinia enterocolitica.* J Gen Microbiol, 1993. **139**(9): p. 2159-65.

52.	Murray, G.L., et al., *Leptospira interrogans requires a functional heme oxygenase to scavenge iron from hemoglobin.* Microbes Infect, 2008. **10**(7): p. 791-7.

53.	Pigeon, C., et al., *A new mouse liver-specific gene, encoding a protein homologous to human antimicrobial peptide hepcidin, is overexpressed during iron overload.* J Biol Chem, 2001. **276**(11): p. 7811-9.

54.	Barton, J.C., C.Q. Edwards, and R.T. Acton, *HFE gene: Structure, function, mutations, and associated iron abnormalities.* Gene, 2015. **574**(2): p. 179-92.

55.	Vert, G., et al., *IRT1, an Arabidopsis transporter essential for iron uptake from the soil and for plant growth.* Plant Cell, 2002. **14**(6): p. 1223-33.

56.	de Lorenzo, V., et al., *Aerobactin biosynthesis and transport genes of plasmid ColV-K30 in Escherichia coli K-12.* J Bacteriol, 1986. **165**(2): p. 570-8.

57.	Bonnah, R.A. and A.B. Schryvers, *Preparation and characterization of Neisseria meningitidis mutants deficient in production of the human lactoferrin-binding proteins LbpA and LbpB.* J Bacteriol, 1998. **180**(12): p. 3080-90.

58.	Allard, K.A., et al., *Purification of Legiobactin and importance of this siderophore in lung infection by Legionella pneumophila.* Infect Immun, 2009. **77**(7): p. 2887-95.

59.	Chatfield, C.H., et al., *The major facilitator superfamily-type protein LbtC promotes the utilization of the legiobactin siderophore by Legionella pneumophila.* Microbiology (Reading), 2012. **158**(Pt 3): p. 721-735.

60.	Chatfield, C.H., et al., *Legionella pneumophila LbtU acts as a novel, TonB-independent receptor for the legiobactin siderophore.* J Bacteriol, 2011. **193**(7): p. 1563-75.

61.	Poole, K., et al., *Multiple antibiotic resistance in Pseudomonas aeruginosa: evidence for involvement of an efflux operon.* J Bacteriol, 1993. **175**(22): p. 7363-72.

62.	Serino, L., et al., *Structural genes for salicylate biosynthesis from chorismate in Pseudomonas aeruginosa.* Mol Gen Genet, 1995. **249**(2): p. 217-28.

63.	Gasser, V., et al., *Catechol siderophores repress the pyochelin pathway and activate the enterobactin pathway in Pseudomonas aeruginosa: an opportunity for siderophore-antibiotic conjugates development.* Environ Microbiol, 2016. **18**(3): p. 819-32.

64.	Koster, M., et al., *Role for the outer membrane ferric siderophore receptor PupB in signal transduction across the bacterial cell envelope.* Embo j, 1994. **13**(12): p. 2805-13.

65.	Tanabe, T., et al., *Identification and characterization of genes required for biosynthesis and transport of the siderophore vibrioferrin in Vibrio parahaemolyticus.* J Bacteriol, 2003. **185**(23): p. 6938-49.

66.	Powell, N.B., et al., *Differential binding of apo and holo human transferrin to meningococci and co-localisation of the transferrin-binding proteins (TbpA and TbpB).* J Med Microbiol, 1998. **47**(3): p. 257-64.

67.	Chen, J., et al., *Transferrin-directed internalization and cycling of transferrin receptor 2.* Traffic, 2009. **10**(10): p. 1488-501.

68.	Enns, C.A., E.A. Rutledge, and A.M. Williams, *The transferrin receptor*, in *Biomembranes: A Multi-Volume Treatise*, A.G. Lee, Editor. 1996, JAIa. p. 255-287.

69.	Bleuel, C., et al., *TolC is involved in enterobactin efflux across the outer membrane of Escherichia coli.* J Bacteriol, 2005. **187**(19): p. 6701-7.

70.	Geoffroy, V.A., J.D. Fetherston, and R.D. Perry, *Yersinia pestis YbtU and YbtT are involved in synthesis of the siderophore yersiniabactin but have different effects on regulation.* Infect Immun, 2000. **68**(8): p. 4452-61.

71.	Bobrov, A.G., et al., *Zinc transporters YbtX and ZnuABC are required for the virulence of Yersinia pestis in bubonic and pneumonic plague in mice.* Metallomics, 2017. **9**(6): p. 757-772.

72.	Waters, B.M., et al., *Mutations in Arabidopsis yellow stripe-like1 and yellow stripe-like3 reveal their roles in metal ion homeostasis and loading of metal ions in seeds.* Plant Physiol, 2006. **141**(4): p. 1446-58.

73.	Divol, F., et al., *The Arabidopsis YELLOW STRIPE LIKE4 and 6 transporters control iron release from the chloroplast.* Plant Cell, 2013. **25**(3): p. 1040-55.

74.	Lill, R., *Function and biogenesis of iron-sulphur proteins.* Nature, 2009. **460**(7257): p. 831-8.

75.	Aw, Y.T.V., et al., *A key cytosolic iron-sulfur cluster synthesis protein localizes to the mitochondrion of Toxoplasma gondii.* Mol Microbiol, 2021. **115**(5): p. 968-985.

76.	Charan, M., et al., *Sulfur Mobilization for Fe-S Cluster Assembly by the Essential SUF Pathway in the Plasmodium falciparum Apicoplast and Its Inhibition.* Antimicrobial Agents and Chemotherapy, 2014. **58**(6): p. 3389-3398.

77.	Ellis, K.E., et al., *Nifs and Sufs in malaria.* Mol Microbiol, 2001. **41**(5): p. 973-81.

78.	Pamukcu, S., et al., *Differential contribution of two organelles of endosymbiotic origin to iron-sulfur cluster synthesis and overall fitness in Toxoplasma.* PLoS Pathog, 2021. **17**(11): p. e1010096.

79.	Bergmann, A., et al., *Toxoplasma gondii requires its plant-like heme biosynthesis pathway for infection.* PLoS Pathog, 2020. **16**(5): p. e1008499.

80.	Harding, C.R., et al., *Genetic screens reveal a central role for heme metabolism in artemisinin susceptibility.* Nature Communications, 2020. **11**(1): p. 4813.

81.	Banerjee, S., et al., *Iron-dependent RNA-binding activity of Mycobacterium tuberculosis aconitase.* J Bacteriol, 2007. **189**(11): p. 4046-52.

82.	Johnson, N.B., et al., *A synergistic role of IRP1 and FBXL5 proteins in coordinating iron metabolism during cell proliferation.* J Biol Chem, 2017. **292**(38): p. 15976-15989.

83.	Hanson, E.S., M.L. Rawlins, and E.A. Leibold, *Oxygen and iron regulation of iron regulatory protein 2.* J Biol Chem, 2003. **278**(41): p. 40337-42.

84.	Hsu, H.M., et al., *Transcriptional regulation of an iron-inducible gene by differential and alternate promoter entries of multiple Myb proteins in the protozoan parasite Trichomonas vaginalis.* Eukaryot Cell, 2009. **8**(3): p. 362-72.

85.	Waldman, B.S., et al., *Identification of a Master Regulator of Differentiation in Toxoplasma.* Cell, 2020. **180**(2): p. 359-372.e16.

86.	Hsu, H.M., et al., *Regulation of nuclear translocation of the Myb1 transcription factor by TvCyclophilin 1 in the protozoan parasite Trichomonas vaginalis.* J Biol Chem, 2014. **289**(27): p. 19120-36.

87.	Yamaguchi-Iwai, Y., A. Dancis, and R.D. Klausner, *AFT1: a mediator of iron regulated transcriptional control in Saccharomyces cerevisiae.* Embo j, 1995. **14**(6): p. 1231-9.

88.	Martínez-Pastor, M.T., A. Perea-García, and S. Puig, *Mechanisms of iron sensing and regulation in the yeast Saccharomyces cerevisiae.* World J Microbiol Biotechnol, 2017. **33**(4): p. 75.

89.	Little, A.S., et al., *Pseudomonas aeruginosa AlgR Phosphorylation Status Differentially Regulates Pyocyanin and Pyoverdine Production.* mBio, 2018. **9**(1).

90.	Brumbarova, T. and P. Bauer, *Iron-mediated control of the basic helix-loop-helix protein FER, a regulator of iron uptake in tomato.* Plant Physiol, 2005. **137**(3): p. 1018-26.

91.	Yuan, Y., et al., *FIT interacts with AtbHLH38 and AtbHLH39 in regulating iron uptake gene expression for iron homeostasis in Arabidopsis.* Cell Res, 2008. **18**(3): p. 385-97.

92.	Beare, P.A., et al., *Siderophore-mediated cell signalling in Pseudomonas aeruginosa: divergent pathways regulate virulence factor production and siderophore receptor synthesis.* Mol Microbiol, 2003. **47**(1): p. 195-207.

93.	Bagg, A. and J.B. Neilands, *Ferric uptake regulation protein acts as a repressor, employing iron (II) as a cofactor to bind the operator of an iron transport operon in Escherichia coli.* Biochemistry, 1987. **26**(17): p. 5471-7.

94.	Koster, M., et al., *Identification and characterization of the pupB gene encoding an inducible ferric-pseudobactin receptor of Pseudomonas putida WCS358.* Mol Microbiol, 1993. **8**(3): p. 591-601.

95.	Zhou, L.W., H. Haas, and G.A. Marzluf, *Isolation and characterization of a new gene, sre, which encodes a GATA-type regulatory protein that controls iron transport in Neurospora crassa.* Mol Gen Genet, 1998. **259**(5): p. 532-40.

96.	Haas, H., et al., *The Aspergillus nidulans GATA factor SREA is involved in regulation of siderophore biosynthesis and control of iron uptake.* J Biol Chem, 1999. **274**(8): p. 4613-9.

97.	Voisard, C., et al., *urbs1, a gene regulating siderophore biosynthesis in Ustilago maydis, encodes a protein similar to the erythroid transcription factor GATA-1.* Mol Cell Biol, 1993. **13**(11): p. 7091-100.

98.	Tran, N.T., et al., *Antiplatelet activity of deferiprone through cyclooxygenase-1 inhibition.* Platelets, 2020. **31**(4): p. 505-512.

99.	Semenza, G.L., *Hypoxia-inducible factor 1: regulator of mitochondrial metabolism and mediator of ischemic preconditioning.* Biochim Biophys Acta, 2011. **1813**(7): p. 1263-8.

100.	Alderton, W.K., C.E. Cooper, and R.G. Knowles, *Nitric oxide synthases: structure, function and inhibition.* Biochem J, 2001. **357**(Pt 3): p. 593-615.

101.	Morishima, Y., et al., *Heme-dependent activation of neuronal nitric oxide synthase by cytosol is due to an Hsp70-dependent, thioredoxin-mediated thiol-disulfide interchange in the heme/substrate binding cleft.* Biochemistry, 2011. **50**(33): p. 7146-56.

102.	Huet, D., et al., *Identification of cryptic subunits from an apicomplexan ATP synthase.* Elife, 2018. **7**.

103.	Mühleip, A., et al., *ATP synthase hexamer assemblies shape cristae of Toxoplasma mitochondria.* Nat Commun, 2021. **12**(1): p. 120.

104.	Salunke, R., et al., *Highly diverged novel subunit composition of apicomplexan F-type ATP synthase identified from Toxoplasma gondii.* PLoS Biol, 2018. **16**(7): p. e2006128.

105.	Seidi, A., et al., *Elucidating the mitochondrial proteome of Toxoplasma gondii reveals the presence of a divergent cytochrome c oxidase.* Elife, 2018. **7**.

106.	Sibley, L.D., R. Lawson, and E. Weidner, *Superoxide dismutase and catalase in Toxoplasma gondii.* Mol Biochem Parasitol, 1986. **19**(1): p. 83-7.

107.	Zhang, X., et al., *Functional characterization of a unique cytochrome P450 in Toxoplasma gondii.* Oncotarget, 2017. **8**(70): p. 115079-115088.

108.	Odberg-Ferragut, C., et al., *Molecular cloning, expression analysis and iron metal cofactor characterisation of a superoxide dismutase from Toxoplasma gondii.* Mol Biochem Parasitol, 2000. **106**(1): p. 121-9.

109.	Brydges, S.D. and V.B. Carruthers, *Mutation of an unusual mitochondrial targeting sequence of SODB2 produces multiple targeting fates in Toxoplasma gondii.* Journal of Cell Science, 2003. **116**(22): p. 4675.

110.	McKie, A.T., et al., *An iron-regulated ferric reductase associated with the absorption of dietary iron.* Science, 2001. **291**(5509): p. 1755-9.

111.	De Silva, D.M., et al., *The FET3 gene product required for high affinity iron transport in yeast is a cell surface ferroxidase.* J Biol Chem, 1995. **270**(3): p. 1098-101.

112.	Ohgami, R.S., et al., *Identification of a ferrireductase required for efficient transferrin-dependent iron uptake in erythroid cells.* Nat Genet, 2005. **37**(11): p. 1264-9.

113.	Levy, J.E., L.K. Montross, and N.C. Andrews, *Genes that modify the hemochromatosis phenotype in mice.* J Clin Invest, 2000. **105**(9): p. 1209-16.

114.	Dupic, F., et al., *Duodenal mRNA expression of iron related genes in response to iron loading and iron deficiency in four strains of mice.* Gut, 2002. **51**(5): p. 648-53.

115.	Immenschuh, S., E. Baumgart-Vogt, and S. Mueller, *Heme oxygenase-1 and iron in liver inflammation: a complex alliance.* Curr Drug Targets, 2010. **11**(12): p. 1541-50.

116.	Ohgami, R.S., et al., *The Steap proteins are metalloreductases.* Blood, 2006. **108**(4): p. 1388-94.
